# Nature of epigenetic aging from a single-cell perspective

**DOI:** 10.1101/2022.09.26.509592

**Authors:** Andrei E. Tarkhov, Thomas Lindstrom-Vautrin, Sirui Zhang, Kejun Ying, Mahdi Moqri, Bohan Zhang, Alexander Tyshkovskiy, Orr Levy, Vadim N. Gladyshev

## Abstract

Age-related changes in DNA methylation (DNAm) form the basis for the development of most robust predictors of age, epigenetic clocks, but a clear mechanistic basis for exactly what part of the aging process they quantify is lacking. Here, to clarify the nature of epigenetic aging, we juxtapose the aging dynamics of tissue and single-cell DNAm (scDNAm) with scDNAm changes during early development, and corroborate our analyses with a single-cell RNAseq analysis within the same multi-omics dataset. We show that epigenetic aging involves co-regulated changes, but it is dominated by the stochastic component, and this agrees with transcriptional coordination patterns. We further support the finding of stochastic epigenetic aging by direct tissue and single-cell DNAm analyses and modeling of aging DNAm trajectories with a stochastic process akin to radiocarbon decay. Finally, we describe a single-cell algorithm for the identification of co-regulated and stochastic CpG clusters showing consistent transcriptomic coordination patterns, providing new opportunities for targeting aging and evaluating longevity interventions.

## Introduction

Epigenetic clocks have been used for almost a decade to accurately predict chronological ages of cells, tissues and organisms^1–6^. More recently, new epigenetic clocks have been introduced that are trained on mortality and/or phenotypic, pathological and physiological readouts^7,8^, as well as on the pace of aging^9^. In all these cases, the basic feature for age prediction or model construction is DNA methylation (DNAm) levels averaged over macroscopic tissue samples, i.e. tissue DNAm levels. The inherent challenge with such an approach is that the DNAm signal is averaged over a large number of different cells present in the tissue. Even though the role of various factors contributing to tissue DNAm changes has been extensively discussed^10–14^, epigenetic aging clocks are still lacking an exhaustive mechanistic explanation. The observed internal age-related changes across all cells could be confounded by multiple factors, such as changes in cell-type composition^15–17^ and errors in DNAm maintenance during cell division and clonal expansion^18–22^. An important advance in this area is the development of a single-cell DNA methylation (scDNAm) clock known as scAge^23^, relying on tissue DNAm data for calibration. Yet, purely clock-based approaches remain phenomenological and do not provide a mechanistic explanation for epigenetic aging.

To clarify the principles and biological mechanisms behind epigenetic aging, we turned our attention to single-cell DNAm and embryonic development data. Using mice as a model, we examined patterns of DNAm across different cells and animals with high inter-cell correlation and used them as a marker of co-regulation. We were able to classify aging changes into two distinct categories: stochastic and co-regulated. Stochastic changes did not form any regular DNAm pattern recurrent across cells and organisms, thus most likely representing the accumulation of epigenetic “rust”. On the other hand, co-regulated changes formed coherent DNAm patterns across different cells and animals, thereby implying the presence of a shared biological mechanism regulating those regions. By chance, random molecular events could also leave similar epigenetic fingerprints in a few cells, but the probability of such an event happening simultaneously in multiple cells of different organisms would be negligible. Conceptually similar to an evolutionary conserved DNA region protected from changes across species, the only possibility of a co-regulated CpG cluster to change its DNAm levels is to do so concordantly within the cell following its DNAm pattern. The co-regulated clusters are thus good candidates for programmatic, regulatory epigenetic changes.

To stress the distinction between stochastic and co-regulated clusters, we run tissue and single-cell DNAm analyses and modeling of aging DNAm trajectories with a stochastic process, along with a supportive single-cell transcriptomic coordination analysis. We start by showing that aging tissue DNAm changes are widespread, but are relatively slow and small in amplitude, with DNAm levels trending towards intermediate values and showing increased heterogeneity with age, in agreement with earlier works studying epigenetic drift^20,24–27^. By considering dominant types of DNAm changes, we find that aging manifests in the exponential decay-like loss or gain of methylation with typical rates, largely independent of the initial level of DNAm^28^. We support our analysis by direct simulation of aging DNAm trajectories within a stochastic model akin to radiocarbon decay. To contrast co-regulation and stochastic patterns during aging, we analyze embryonic development scDNAm data as a control for co-regulated changes because of a tightly controlled genetic program that governs development. Finally, in scRNAseq data, we find supportive evidence of significant global coordination levels (GCL)^29,30^ of gene expression for genes associated with co-regulated CpG clusters compared to non-significant GCL for stochastic CpG clusters.

The current approaches for building epigenetic clocks are mostly based on penalized regression (Lasso, Ridge or ElasticNet), which penalize correlation across CpG sites in order to reduce the number of CpG sites used to build the clock. Therefore, the penalization would suppress the contribution of co-regulated clusters because of their high inter-cell correlation. The predictions of age or other traits would be insensitive to the composition of the clock in the correlative analysis. Thus, the currently built clocks may be biased towards stochastic CpG sites. Though, since clocks typically comprise a few hundred CpGs, in addition to stochastic CpGs, the same penalization may select (but is not guaranteed to do so) a few CpG sites per each of multiple aging-related pathways^13^, thus effectively amplifying the aging trend in methylation changes by averaging out noise proportionally to the square root of a number of CpGs as defined by the standard deviation of a sample mean. Hence, a clock with a hundred CpG sites is enough to suppress the noise in aging signal by a factor of 10, which may explain why such epigenetic clocks exhibit high robustness and precision in age prediction (*r*^2^ > 0. 8 on an independent validation dataset^31^). However, different parts of co-regulated clusters and stochastic sites may respond qualitatively differently in the case of interventions applied to an organism. A more mechanistic understanding of epigenetic aging based on co-regulation may improve performance of epigenetic clocks in evaluating anti-aging interventions. The current approaches for building epigenetic clocks mix up two qualitatively different components (co-regulated and stochastic) that therefore limit their predictive power for testing interventions.

Summarizing the multifaceted evidence emerging from our analyses, aging scDNAm changes are dominated by the stochastic component, in agreement with the concept of aging as the entropic loss of complexity^32,33^. We corroborate our finding with supportive transcriptional coordination patterns in scRNAseq data matching scDNAm patterns. The stochastic component of aging^34^ may be a good indicator of cumulative damage arising from internal processes together with the deleteriousness of the environment in which an organism lives. Conversely, co-regulated CpG clusters, despite their sparsity, may show promise as better candidates for capturing the effects of target-specific anti-aging interventions.

## Results

### Tissue epigenetic aging

To assess the behavior of DNAm over the time course of mouse life, we divided it into three intervals according to the typical aging dynamics of physiological readouts: development, functional aging and multimorbidity^35,36^ (**Fig. 1a**). Similar life-course staging can be applied to humans^37–39^. Development, especially early development, is known to be a tightly regulated, programmed process, even if molecular aging already exists there^40^. Functional aging corresponds to the period of life between the end of development, accompanied by a gradual functional decline without major comorbidities. Multimorbidity is the final stage of life, when the functional decline becomes incompatible with organ and tissue function and organismal survival. We first focused our analyses on the period of functional aging and then made use of developmental data for uncovering dissimilarities in DNAm dynamics between these two life stages.

**Fig. 1.**
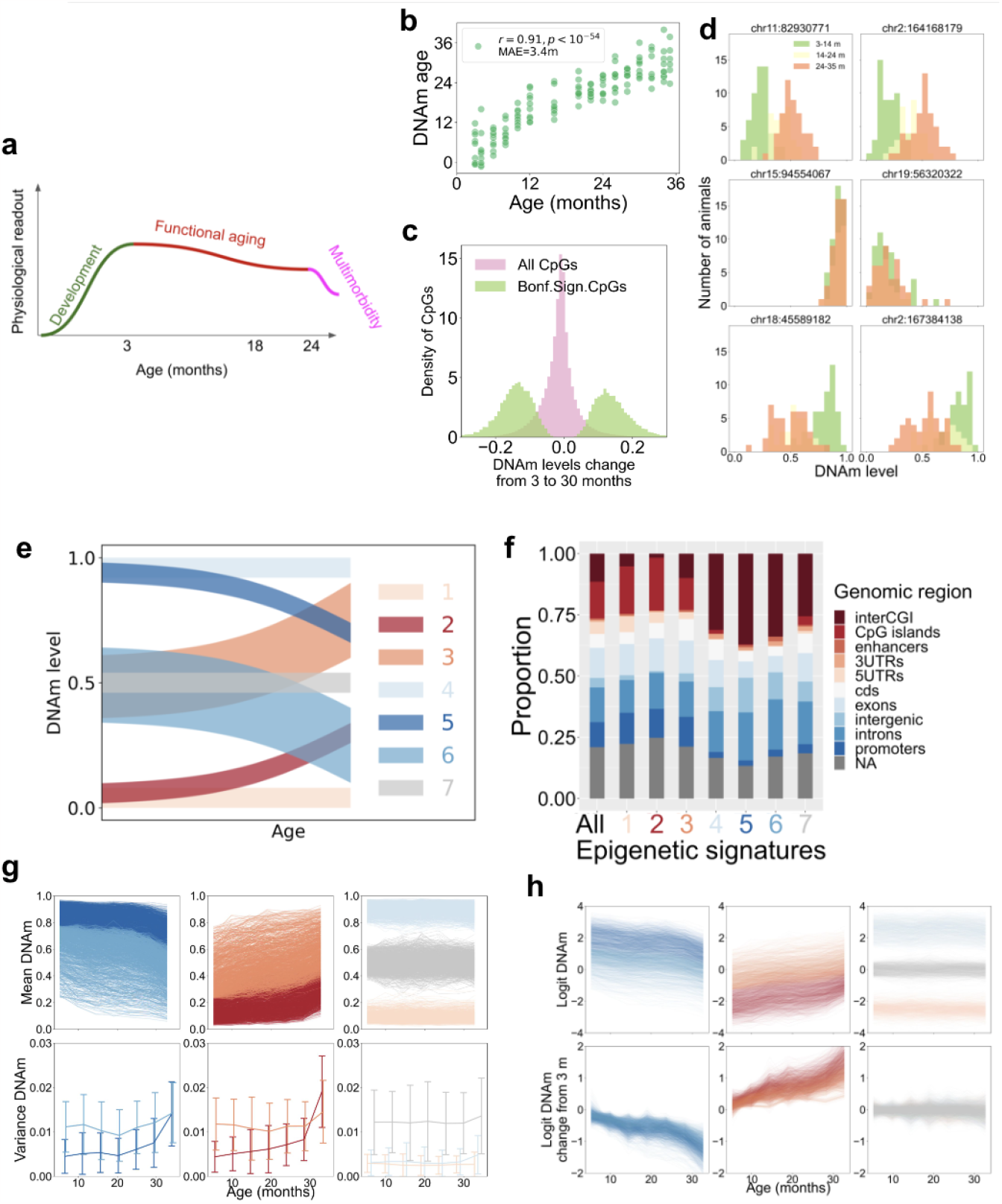
Key features of tissue DNAm changes during aging. **a**. Three major life stages of mice represented by age-related changes in function — development, functional aging and multimorbidity^35,36^. **b**. Predicted DNAm age of animals (16 age groups, from 3 till 35 months old) based on the Petkovich *et at*. dataset^4^. **c**. The range of DNAm changes at CpGs sites in the dataset. Histograms are shown for all CpG sites and for the CpGs significantly changing with age. **d**. Six representative CpG histograms in aging mouse cohorts of 3-14 (green), 14-24 (yellow) and 24-35 (orange) months old for the following CpG sites: significantly hypermethylated with age (upper row), not changing with age (middle row), and significantly hypomethylated with age (lower row)^4^. **e**. Schematics of seven dominant aging DNAm dynamics observed in experimental data. Other types of dynamics are rare but may also be present in the data. **f**. Genomic enrichment analysis for the CpG sites comprising the seven aging dynamics from **e**. Enrichment profiles are similar for the CpG sites sharing the direction of age-related changes (dynamics 2 and 3, and 5 and 6). Note that entropic changes do not share a common genomic profile (2 and 5), and the same applies to those decreasing entropy(3 and 6). **g**. Aging dynamics of mean DNAm levels (upper row) and the variance of DNAm levels (lower row) for the seven types of aging dynamics from **e**. Color scheme is the same as in **e. h**. Aging dynamics of logit of mean DNAm levels (upper row), and of the logit of mean DNAm levels relative to the logit of mean DNAm levels at 3 months of age. The logit was defined as 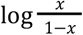, where *x* is the DNAm level.

We first analyzed total blood (bulk) DNA methylation (DNAm) changes during aging of male C57Bl/6 mice (16 age groups from 3 to 35 months, 8 animals per group) based on our previous reduced representation bisulfite sequencing dataset^4,41^. There were 268,044 CpG sites that significantly correlated with age (13.6% out of all 1,976,056 CpG sites measured in at least one sample with sequencing depth 10X, and after the Bonferroni correction for multiple testing, 16,889 CpGs (0.85% of all sites). Given that a substantial fraction of CpGs change significantly with age, a straightforward prediction of chronological age based on the DNAm levels and elastic net regression could be made (**Fig. 1b**), as was previously done in the case of other aging clocks. Epigenetic clocks typically comprise a few hundred of CpG sites out of hundreds of thousands that significantly change with age, hence the major challenge is how to select CpGs that contain the most biologically relevant information about aging and would produce valuable biomarkers of this process; for example, one may use additional information, such as genetic data^42^, for this purpose. Interestingly, the overall change of DNAm level for each of the numerous age-related CpG sites over the period of functional aging was small. The largest change of methylation was ∼30%, whereas for most of the CpG sites it was below 10% throughout lifespan (**Fig. 1c**). In some cases, even a small (e.g. 10%) change in DNAm level may result in significant consequences for cells (e.g., activation of retrotransposons or pro-apoptotic genes) and cause health outcomes for the organism in the long-term. Nevertheless, on average, we observed global DNAm changes in the genome that only slightly change their normal DNAm level throughout lifespan. For illustration, we present six representative CpG histograms of aging mouse cohorts for pairs of CpGs significantly hypermethylated with age, not changing with age, and significantly hypomethylated with age (**Fig. 1d**).

We further classified the aging trajectories of DNAm into seven categories depending on the initial methylation value at the end of development (at 3 months), and the direction of subsequent changes (**Fig. 1e, g, h**). Dynamics 1, 4 and 7 represent CpGs, whose methylation levels remain constant with age and differ only by the initial DNAm level. Dynamics 2 and 3 correspond to the growth of methylation with age, and dynamics 5 and 6 to the loss of methylation. Dynamics 2 and 5 trend towards the methylation level of 0.5 according to the growth of entropy. At the same time, dynamics 3 and 6 correspond to an apparent loss of entropy. To characterize these dynamics, we carried out genomic enrichment analyses of epigenetic signatures corresponding to the seven types of dynamics (**Fig. 1f**). Dynamics 2 and 3 (increasing methylation) and dynamics 5 and 6 (losing methylation) clustered together, whereas entropic dynamics 2 and 5 or the ones decreasing entropy, 3 and 6, ended up in different clusters, thus supporting the existence of two distinct processes driving methylation levels up and down independent of the initial methylation. Thus, aging dynamics cluster according to the direction of DNAm level change over functional aging rather than according to the initial methylation level at the completion of development. The dynamics corresponding to the loss of methylation and higher initial methylation levels (4, 5, 6 and 7) compared to those gaining methylation or having low initial methylation levels (1, 2 and 3) are enriched with inter-CpG island regions, or open sea, introns, intergenic regions and 3’-UTRs, i.e. generally non-functional regions, in agreement with prior studies^20,25,26^. Likewise, they are depleted for CpG islands, coding sequences, exons, 5’-UTRs and promoters.

Interestingly, the aging change was accompanied by the growth of heterogeneity of DNAm levels within cohorts of the same chronological age, which is indicated by the broadening of methylation level distribution with age (**Fig. 1d)**, and the growth of DNAm level variance (**Fig. 1g**). Not only the mean DNAm level for CpGs corresponding to dynamics 2 and 5 showed a pronounced change with age, but also the heterogeneity of DNAm levels in age cohorts grew with age (**Fig. 1g**). Additionally, they shared a similar rate of DNAm change notwithstanding the initial methylation level. We applied the logit transformation of DNAm levels, 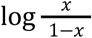(x is the DNAm level) and found that after this transformation the changes were essentially linear (**Fig. 1h**), thus resembling exponential-decay-like changes in the average methylation levels within the defined dynamics of DNAm changes. This is consistent with the aging-related increase in entropy metrics based on various molecular and functional models and readouts^12,41,43–46^.

The above analysis of blood DNAm aging changes can be summarized by the following key points. DNAm changes during functional aging are omnipresent in the genome, but are relatively slow and small in the amplitude. DNAm levels, in general, trend towards the methylation level 0.5 and display increased heterogeneity with age, but this involves different mechanics. Aging DNAm dynamics can be clustered into seven dominant types. Aging manifests in the exponential-decay-like loss (or gain) of methylation with typical rates, largely independent of the initial level of DNAm. These key points suggest that the DNAm aging changes demonstrate common features indicative of a stochastic process.

### Stochastic single-cell model for simulating tissue epigenetic aging

To explain a part of age-related DNAm changes caused by stochastic damage accumulation, we tested the hypothesis that it is possible to reproduce experimental DNAm aging trajectories with a stochastic single-cell model (see Methods in Supplementary Information). For simplicity, the stochastic model considers only a single allele per CpG and assumes that the DNAm dynamics for a CpG site in a cell are purely stochastic and controlled by three parameters: *q* — initial methylation level, *p* _*d*_ and *p*_*m*_ — rates of demethylation and methylation in a unit of time. We illustrate the stochastic model with an example of two CpG sites in a tissue sample of 40 cells — for CpG 1 the rate of methylation is higher than the rate of demethylation, whereas the initial level of methylation is low, and for CpG 2 the parameters are the opposite (**Fig. 2a**). As time passes, some of the cells would flip their methylation state according to the predefined probabilities. However, the tissue DNAm would average single-cell levels of methylation across all cells and produce a single number for the mean DNAm level, which is predicted by the model. For a sample of *n* cells, the CpG site is methylated in *n*_*m*_ cells. During functional aging, the CpG can randomly acquire or lose methylation according to the probability rates *p* _*d*_ and *p*_*m*_ defined above. The model predicts the equation for the aging trajectory of the average level of methylation across the whole sample *x*(*t*) = *n*_*m*_ (*t*)/*n* to be an exponential curve (**Fig. 2b**):

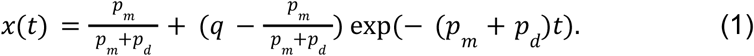

To compare real aging trajectories of CpG sites with the model predictions, we analyzed the DNAm aging dynamics^4^ and calculated the rates *p*_*m*_, *p* _*d*_ for all sites changing significantly with age (**Fig. 2c**), and plotted the aging dynamics for 90 CpGs comprising the Petkovich *et al*. epigenetic clock (**Fig. 2d, left**). Next, we fitted the experimental DNAm levels dynamics to *x*(*t*) for each clock CpG site (**Fig. 2d, middle**). We also plotted a subset of trajectories corresponding to the parameters *q, p* _*d*_, *p*_*m*_ randomly drawn from the uniform distributions on the following intervals: *q* ∈ [0, 1], *p*_*m*_, *p* _*d*_ ∈ [0, 0. 045] (**Fig. 2c**) for each CpG site (**Fig. 2d, right**).

**Fig. 2.**
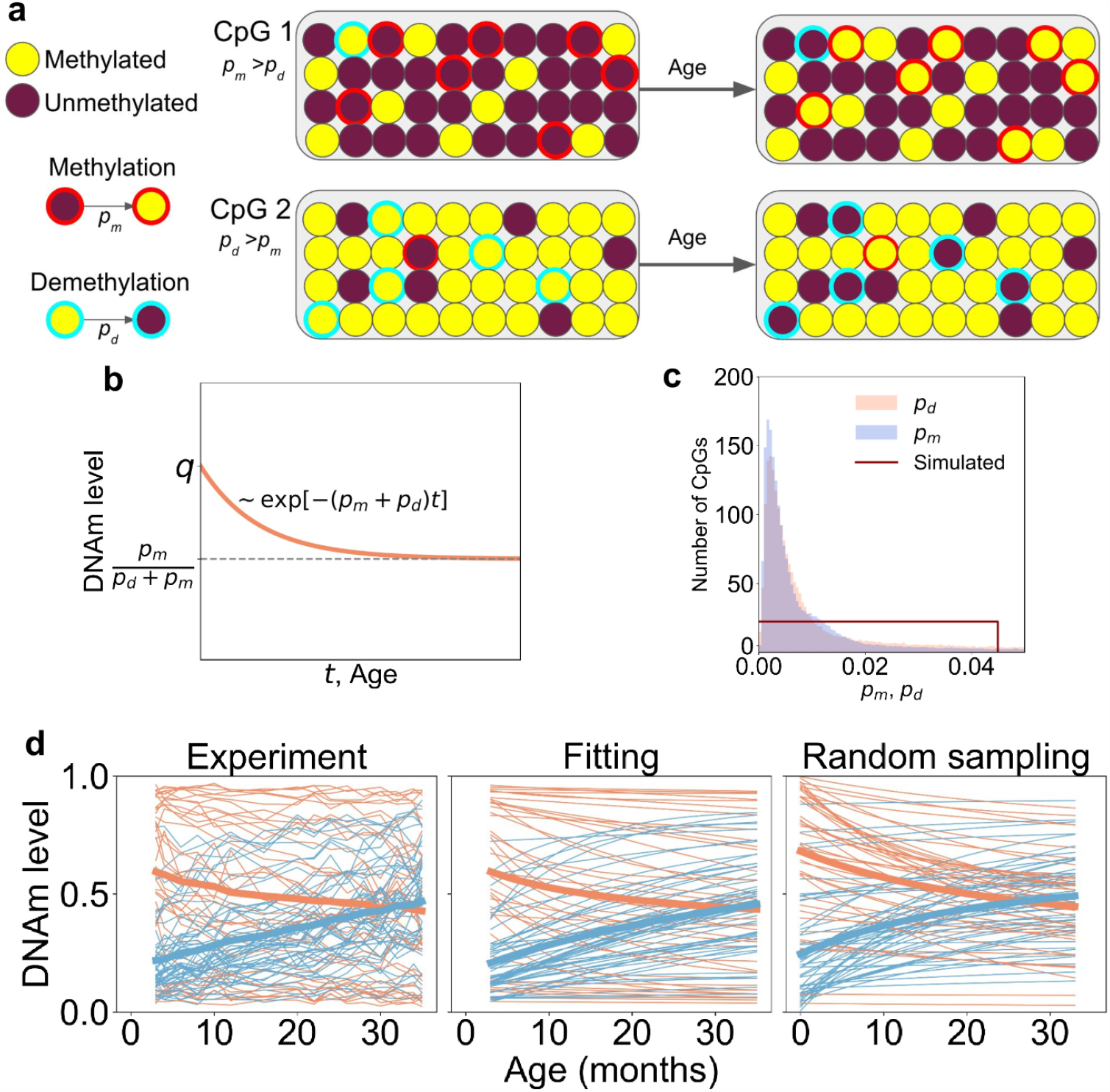
Stochastic single-cell model for modeling tissue DNAm dynamics. **a**. Sketch for a three-parameter stochastic decay model: *q* — initial methylation level, *p* _*d*_ and *p*_*m*_ — rates of demethylation and methylation in a unit of time. The flips of methylation occur randomly with the corresponding rates. The behavior of two CpG sites in single cells comprising a tissue sample is illustrated. CpG 1 is initially hypomethylated and gets more methylated with age, whereas CpG 2 is initially hypermethylated and loses its methylation. **b**. The stochastic decay model produces exponential decay law for the dynamics of average methylation. The rate of decay equals the sum of *p*_*m*_ and *p* _*d*_ . The limiting value is defined by the dynamic equilibrium of methylation and demethylation. **c**. Distributions of *p*_*m*_ and *p* _*d*_ for the sites significantly changing with age^4^, which were observed in the experiment. Random sampling was performed for a uniform distribution of *q, p*_*m*_ and *p* _*d*_ on the following intervals of parameters: *q* ∈ [0, 1], *p*_*m*_, *p* _*d*_ ∈ [0, 0. 045]. **d**. Dynamics of CpG sites included in the Petkovich et al.^4^ DNAm clock (left), results of fitting of those trajectories to the exponential decay defined in **c** (middle), and random sampling of aging trajectories **b** sampled from the distribution in **c** (right). Blue lines show trajectories gaining methylation, and orange lines — losing methylation. Thick orange and blue lines are the mean trajectories of orange and blue lines.

The stochastic model predictions can fit the observed behavior of clock CpG aging trajectories (**Fig. 2d left** and **middle**). At the same time, the semi-quantitative similarity of experimental and randomly sampled trajectories (**Fig. 2d left** and **right)** suggests that the stochastic model is able, in a very crude approximation, to qualitatively reproduce the experimental behavior within a stochastic framework. The distributions of parameters *q, p*_*d*_, *p*_*m*_ for simulated trajectories can be characterized by a single characteristic parameter — the maximal rate of exponential decay (*p*_*m*_ + *p*_*d*_)_*max*_ (**Fig. 2c**). This parameter may be used for crude characterization of instability of the aging epigenome in order to estimate the characteristic age, when epigenetic instability becomes evident. One can potentially refine such an approximation to take into account more subtle features of the aging epigenome.

### Tissue epigenetic clocks are agnostic of the single cell patterns of aging

Different single-cell DNAm distributions can produce the same tissue DNAm levels. Specifically, single-cell aging changes can be caused by drastically different biological processes without being detected by tissue DNAm signals. To illustrate this point, we sketch three possible scenarios of single-cell DNAm changes that correspond to the same tissue DNAm pattern (**Fig. 3a**). The left half of CpGs were initially unmethylated in all cells and organisms, and the right half — methylated. Then, over the course of aging, each unmethylated CpG site acquires 20% of methylation, and each methylated CpG site loses 20% of methylation. Three single-cell scenarios are shown: stochastic, co-regulated and mixed. The stochastic scenario assumes that all changes are scattered across cells in an uncorrelated manner — the DNAm changes occur independently at CpGs in cells and organisms. The co-regulated scenario represents a model wherein there are two states for a cluster of CpGs, young and old, and during aging 20% of cells switch from the young state to the old state. This scenario corresponds, for instance, to the case of senescent cells, whose population grows with age. The third scenario is a mixture of the two described above — all cells accumulate stochastic changes of methylation, whereas some genomic regions are co-regulated. Tissue DNAm data generated in these three scenarios would indicate the same 20% change of DNAm levels, thus being unable to distinguish these scenarios. In real scDNAm data, low sequence coverage produces a severely sparse signal (**Fig. 3b**), making the analysis of scDNAm extremely challenging but not impossible in mitotic or clonally expanding cells. Below, for our analysis, we use the data on single-cell muscle stem cells and embryonic cells to partially overcome this challenge.

**Fig. 3.**
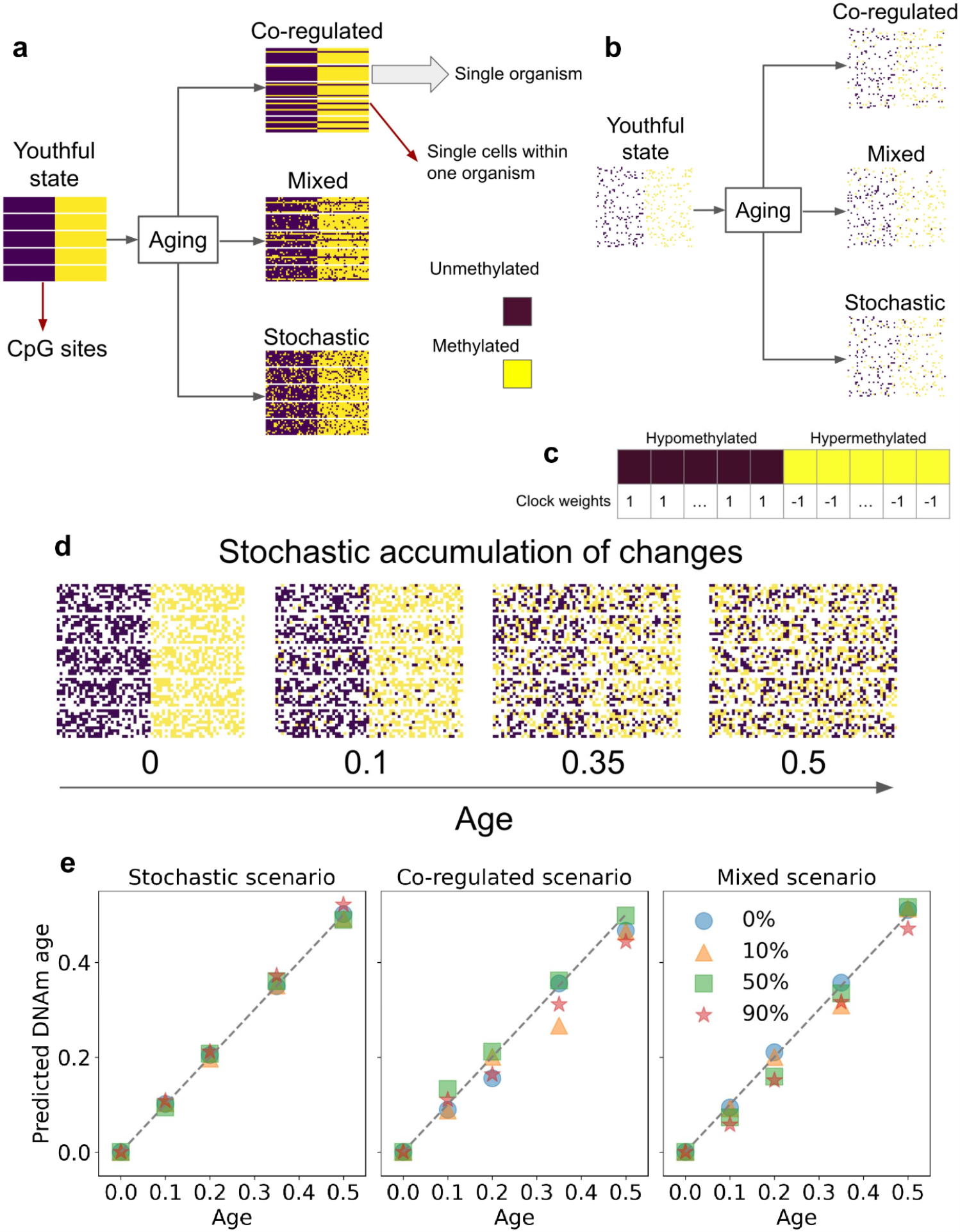
Three single-cell scenarios of DNAm aging changes. **a**. An example of three single-cell scenarios used to produce the same 20% change of tissue DNAm levels with age: co-regulated, stochastic and mixed. The example is shown for 5 aging animals (horizontal rectangles) with DNAm levels measured for a set of cells (represented by rows in the rectangles). Each CpG is represented by a column, the left half of the CpGs are assumed to be hypomethylated in the youthful state, whereas the right half is initially hypermethylated. Co-regulated aging changes occur in a correlated manner within one cell (changes of different CpG sites are correlated), and in a coherent way in different cells and animals. Stochastic changes are uncorrelated among different cells and among different CpG sites. Mixed changes are a combination of the two. Tissue DNAm clocks are unable to distinguish these three scenarios since they all correspond to the same change in tissue DNAm levels. **b**. Realistic coverage of NGS in single cells would produce a sparse subset of CpGs measured in only a handful of cells. The sparsity in the plot is 90%. **c**. A hypothetical clock with the left half of all CpG sites hypomethylated in the youthful state, having weight +1, whereas the right half of CpG sites hypermethylated in the youthful state, having weight -1. For age prediction, the clock weights are normalized by the number of CpGs in the clock. **d**. Illustration of the dynamics of stochastic accumulation of epigenetic changes with age. Age corresponds to the fraction of average DNAm levels (from 0 to 0.5). The sparsity level is 50%. **e**. Prediction of the hypothetical clock from **c** (normalized to have a unit length) on the simulated data for the three scenarios from **a**. The clock from **c** is able to accurately predict chronological age for any used single-cell scenario, and for any NGS coverage (see legend for sparsity levels).

We use a hypothetical example of a clock built for the simulated single-cell scenarios (**Fig. 3c**). Since tissue DNAm changes are the same 20% for all CpGs, the clock has two constant parts: it has weights +1 for the right half of CpGs, and -1 for the left half (normalized to the total number of CpGs). We illustrate the dynamics of aging changes in the stochastic scenario example (**Fig. 3d**). We show that the prediction of chronological age is possible no matter what single-cell scenario or the level of sparsity of the scDNAm signal were used to simulate the aging dynamics (**Fig. 3e**). At the same time, the clock is unable to distinguish the single-cell scenarios used to generate scDNAm levels. The mechanism behind such a clock might be a gradual accumulation of stochastic damage, or the growth of the population of “old” cells, or the mixture of the two. In all cases the predictive power of the clock is based on the inevitable change of average methylation levels with age. Further, we turn to real scDNAm data to clarify what scenario is realized in an experiment.

### Single-cell epigenetic dynamics during functional aging and embryonic development

We analyzed two scDNAm datasets that provide high NGS coverage for each cell: mouse embryos prior and during gastrulation^48^ and aging muscle stem cells (muSC)^49^. In the case of aging muSCs, we had 275 single cells derived from four 2-month-old mice and two 24-month-old mice (one old mouse was censored because of the low coverage of NGS in its cells, see Methods for more detail). First, we filtered CpGs by coverage: each CpG must be measured in at least 15 cells of young mice, and 15 cells of old mice. Out of 35,584,147 CpGs measured in at least one cell, only 155,359 had sufficient coverage and passed the filter. Second, we identified CpGs changing significantly with age, thus leaving 502 CpG sites. Within those CpGs, we managed to observe all seven types of tissue DNAm aging dynamics present in **Fig. 1d** (**Fig. 4a**). For convenient comparison with the tissue DNAm dynamics, we present both raw and compressed scDNAm data, where for each CpG we omit non-measured cells and collapse all cells for the young mice to the top, and for the old mice to the bottom of the panel (**Fig. 4a left** and **right**).

**Fig. 4.**
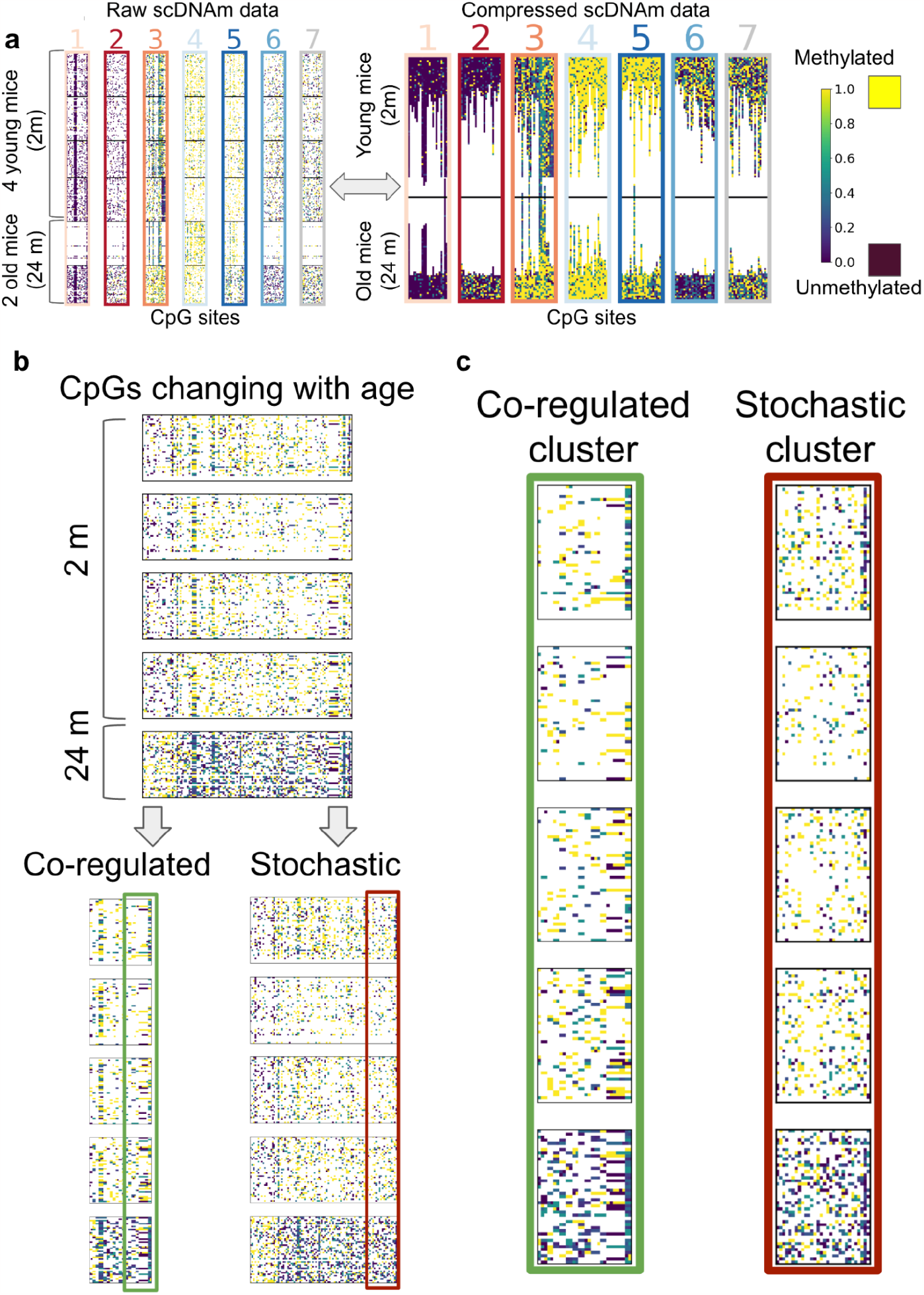
Single-cell DNAm dynamics during functional aging. **a**. All seven types of aging trajectories are present in muscle stem cells (muSC) scDNAm data^47^ (**Fig. 1e**). Raw sparse scDNAm data are shown for a subset of 275 cells from 4 young mice (2 months old) and 2 old mice (24 months old), single cells from which represent horizontal lines (**left panel**). Black lines separate cells corresponding to different mice. For each CpG (vertical column), compression omits non-measured cells, and shows only the measured ones (**right panel**). All measured cells for the young mice are collapsed to the top of the figure, and for the old mice to the bottom. One old mouse showed a significantly lower coverage than other animals and was censored from the following analyses (see Methods for more detail). **b**. Dynamics of a subset of CpG sites that change significantly with age in scDNAm data (from 2-month-old to 24-month-old mice) (**upper panel**). All age-related CpGs were classified into co-regulated and stochastic clusters by their inter-cell correlation (**lower panel**). The co-regulated cluster has characteristic stripes of CpGs whose DNAm levels change concordantly (**lower left panel**). The stochastic cluster does not show strong correlation among different cells (**lower right panel**). The largest co-regulated cluster (**green box**), and a stochastic cluster of the same size (**brown box**) were chosen for an additional close-up view. **c**. Close-up view of the co-regulated cluster (**left**) and a stochastic cluster (**right**) from **b**. The co-regulated cluster indicates the stretches of CpGs that show high correlation among themselves, and change their DNAm levels as a single unit. The stochastic cluster does not have similar clustering of CpG sites. Each CpG site in the stochastic cluster changes independently from others.

Out of 502 CpGs changing methylation with age, we identified 121 CpG sites increasing methylation and 381 CpG sites losing methylation with age (**Fig. 4b**). To identify co-regulated clusters of CpGs, for each pair of CpGs, we calculated the Pearson inter-cell correlation coefficient separately for young mice, old mice, and for all mice together. The values of correlation had to be higher than a given threshold in the range from 0. 1 to 0. 9 in all three cases to include the pair of CpGs into the co-regulated cluster (for illustration purposes, we use threshold correlation value 0. 5, see Methods and **Ext. Data Figs. 2, 5** and **6** for more details on the stability of our analyses to the choice of the threshold, and **Ext. Data Fig. 8** for extensive tests of the algorithm on simulated data). For inter-cell Pearson correlative analysis, we used a custom Python code taking into account only those DNAm values for CpGs that had a non-zero coverage. The CpGs with zero coverage were not used for calculating the Pearson correlation. The method of identifying co-regulated clusters is conceptually similar to the global coordination level (GCL) metric developed independently for scRNAseq data analysis^29,30^. The majority of age-related CpG sites, 92%, changed according to the stochastic scenario, whereas 8% of the CpG sites changed in a co-regulated manner. Such a definition of co-regulation is relatively strict, and would rather tend to mark a CpG site stochastic if it has a low coverage. For a larger number of sequenced cells, the method would be able to identify more co-regulated clusters. Therefore, the ability to identify real co-regulated clusters may be strongly influenced by experimental limitations. With the advances in sequencing technologies, it may be possible to improve the quality of prediction of co-regulation.

Commonly used for building epigenetic clocks, penalized regression methods have the advantage of selecting CpG sites that add the most new information to the regression model at the cost of reducing the number of collinear CpG sites. However, their major drawback in the context of scDNAm data is that they may be biased towards stochastically changing CpG sites (**Ext. Data Fig. 1, Lasso** and **ElasticNet clocks**). The reason for the bias is that the CpG sites comprising co-regulated clusters are strongly collinear to each other. For the purpose of mathematical regression, they do not add any additional information to the model, whereas the stochastic sites are poorly correlated with each other, and the inclusion of each additional stochastic CpG site is advantageous for the algorithm performance. The biological meaning of co-regulation implies that there is a well-defined biological process that controls each co-regulated cluster and does not allow the components of this cluster to independently accumulate epigenetic damage. However, stochasticity may affect a co-regulated cluster as a whole if a damaging event affects an upstream mechanism governing the co-regulation, for example, the DNA regions comprising a co-regulated cluster may be located close to each other in the nucleus or may have similar methyltransferase/demethylase recognition sites. Therefore, the mathematical collinearity is coherent with the biological meaning behind the cluster, which is ignored by penalized regression models.

To test the co-regulation scenario in the context of developmental genetic programs, we analyzed embryonic scDNAm data^48^. There, we had 758 single-cell samples for embryonic days E4.5, E5.5, E6.5 and E7.5 collected from 28 embryos. First, we chose the CpGs changing during functional aging (**Fig. 5a left**) that were also measured in the embryos (370 out of 502 CpGs). These CpGs changed in a coherent co-regulated and program-like manner: at E4.5 the global methylation was low, whereas by E5.5 it was largely set to a methylated state, and only slightly further increased during E6.5 and E7.5 (**Fig. 5a right**). Overall, the dynamics of scDNAm changes during functional aging was different from that during gastrulation. Some CpGs losing methylation in old animals gained it during gastrulation, consistent with the previously observed DNAm age dynamics based on clocks^40^. The global change of DNAm during gastrulation follows a trend opposite to the aging changes of DNAm age. At the same time, co-regulated aging clusters also changed during embryonic development, which may signify the developmental activation of the same biological mechanisms becoming prominent later in life. Moreover, some CpGs that get unmethylated during aging, gain methylation during embryonic development.

**Fig. 5.**
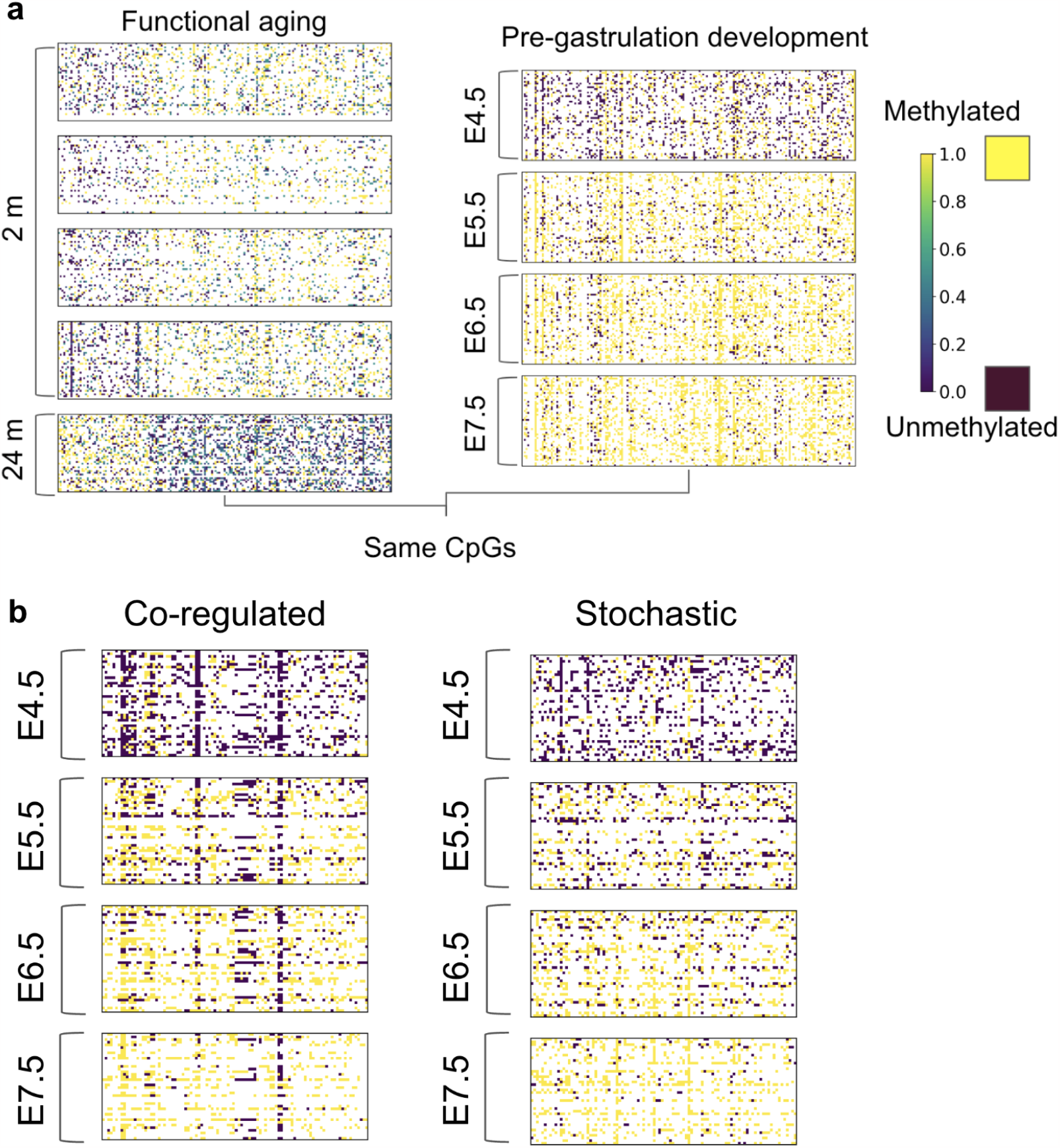
Comparison of single-cell DNAm dynamics during functional aging and embryonic development. **a**. Comparison of aging dynamics of CpGs (**left**) to their dynamics in mouse embryos before and during gastrulation (**right**). Common CpG sites measured in both experiments were used for plotting. Both hyper- and hypo-methylated with age CpGs (**left**) were initially hypomethylated at embryonic day E4.5, and largely acquired global methylation by day E5.5 (**right**). Afterwards little qualitative change of methylation was detected up to day E7.5. The qualitative character of changes during functional aging comprises both co-regulated and stochastic clusters, whereas the pre-gastrulation development is characterized by the program-like global hypermethylation. **b**. Co-regulated and stochastic clusters identified for changes for days E4.5, E5.5, E6.5 and E7.5. The dominant process is gaining methylation at all co-regulated CpG sites.

Two factors complicate our analysis of co-regulation in embryonic datasets. First, the cells were extracted from 28 different embryos that may increase inter-embryo heterogeneity in the scDNAm signal, which would disfavor co-regulation. Second, the global change of methylation during gastrulation represents a complication for our algorithm — in the case of global changes of methylation, the inter-cell correlation turns out to be less meaningful because of minor variation of methylation levels among cells and organisms at each developmental stage. In a hypothetical example of methylation switching from 0 to 1 in all cells of all organisms, it is unclear how to identify individual clusters responsible for the global change of methylation. Therefore, we examined co-regulation in those regions that do not exactly follow the global wave of methylation. To identify such co-regulated clusters in the DNAm changes pre- and during gastrulation, we filtered CpGs by coverage: each CpG must be detected in at least 25 cells for samples corresponding to each embryonic day. Out of 20,073,742 CpGs measured in at least one cell only 44,711 had sufficient coverage and passed the filtering, whereas 6,000 CpGs changed significantly with age. Application of our algorithm with the correlation threshold 0. 3 produced a co-regulated cluster of 191 CpGs, and 5,809 stochastic CpGs (**Fig. 5b**). Due to inter-embryo heterogeneity, a relatively low sequencing coverage and high sequencing noise, in the embryonic dataset, we could identify less than 5% of co-regulated CpGs in the background of the global methylation (**Ext. Data Fig. 4**).

Overall, our analysis showed, with the caveats discussed above, that the major component in the aging scDNAm signal is stochastic. However, some aging CpG sites are consistently identified as co-regulated, with several detected clusters spanning extensive genomic regions. In the following section, we present a biological annotation of the identified clusters.

### Biological annotation of co-regulated and stochastic clusters

The top two co-regulated CpG clusters with the strongest signal (**Fig. 4c)** correspond to the genomic region chr7:7,299,638-7,299,695 and overlaps with the enhancer (chr7:7,299,662-7,299,876) of gene *Clcn4* (chloride voltage-gated channel 4), and the region (chr18:61,035,975-61,035,992) in the enhancer of gene *Cdx1* (caudal type homeobox 1). Variants in gene CLCN4 in humans are associated with a rare X-linked CLCN4-related neurodevelopmental disorder (CLCN4-NDD)^44,45^. Human CDX1 is a tumor suppressor gene, loss of function variants in which are associated with intestinal tumors^46^. Both co-regulated CpG clusters are in the genomic regions containing the marks of high H3K4me3 and H3K27ac indicating active transcription and an open chromatin state^47^. The active transcription of *Clcn4* and *Cdx1* and their clinical importance may be the reason why the CpGs located in this region are strongly co-regulated during aging. At the same time, stochastic CpGs with the lowest signal of co-regulation (for example, chr1:85270111 in *C130026l21Rik*, chr2:128286757 in LncRNA *Morrbid*, chr4:43660058 in *Fam221b*, chr8:115878285 distant from *Maf*, chr9:68760734 close to *Rora*, chr12:116705833 close to *Ptprn2*, chr18:46761319 close to *Ap3s1*) do not form contingent groups and lie far from active promoters/enhancers.

To obtain more CpGs belonging to co-regulated and stochastic clusters for subsequent biological annotation, we modified our filters. First, we lowered the filter for CpGs based on coverage: instead of requiring 15 young and 15 old cells, we required 5 cells in each case. We identified 5,999,943 CpGs out of 35,584,147 CpGs measured in at least one cell, including 51,895 CpGs that changed significantly with age. In this case, the co-regulated and stochastic clusters comprised 2,764 and 49,131 CpGs, respectively. As expected from the algorithm’s properties being biased towards a stricter criterion for co-regulated sites, the fraction of stochastic sites increased from 92% to 94.7%. In addition to 51,895 CpGs changing methylation with age, we selected 51,895 random CpGs from the genome for subsequent enrichment analyses.

We calculated evolutionary conservation scores phyloP^50,51^ and phastCons^52–56^ for co-regulated and stochastic clusters (**Fig. 6a, Ext. Data Fig. 2d**), as well as for genomic regions corresponding to dynamics 2 and 3, hypermethylated with age, and dynamics 5 and 6, hypomethylated with age (**Fig. 6b**). Co-regulated clusters showed a significantly higher evolutionary conservation than stochastic clusters and random regions, in agreement with the hypothesis of a tighter regulatory control and a higher biological importance of those regions (**Fig. 6a**). At the same time, stochastic regions showed a significantly lower evolutionary conservation than random genomic regions (**Fig. 6a**). There was no significant difference among the evolutionary conservation for hypermethylated, hypomethylated and random regions (**Fig. 6b**). The results for evolutionary conservation were stable and consistent for any chosen correlation threshold from 0. 1 to 0. 9 (**Fig. 6c**).

**Fig. 6.**
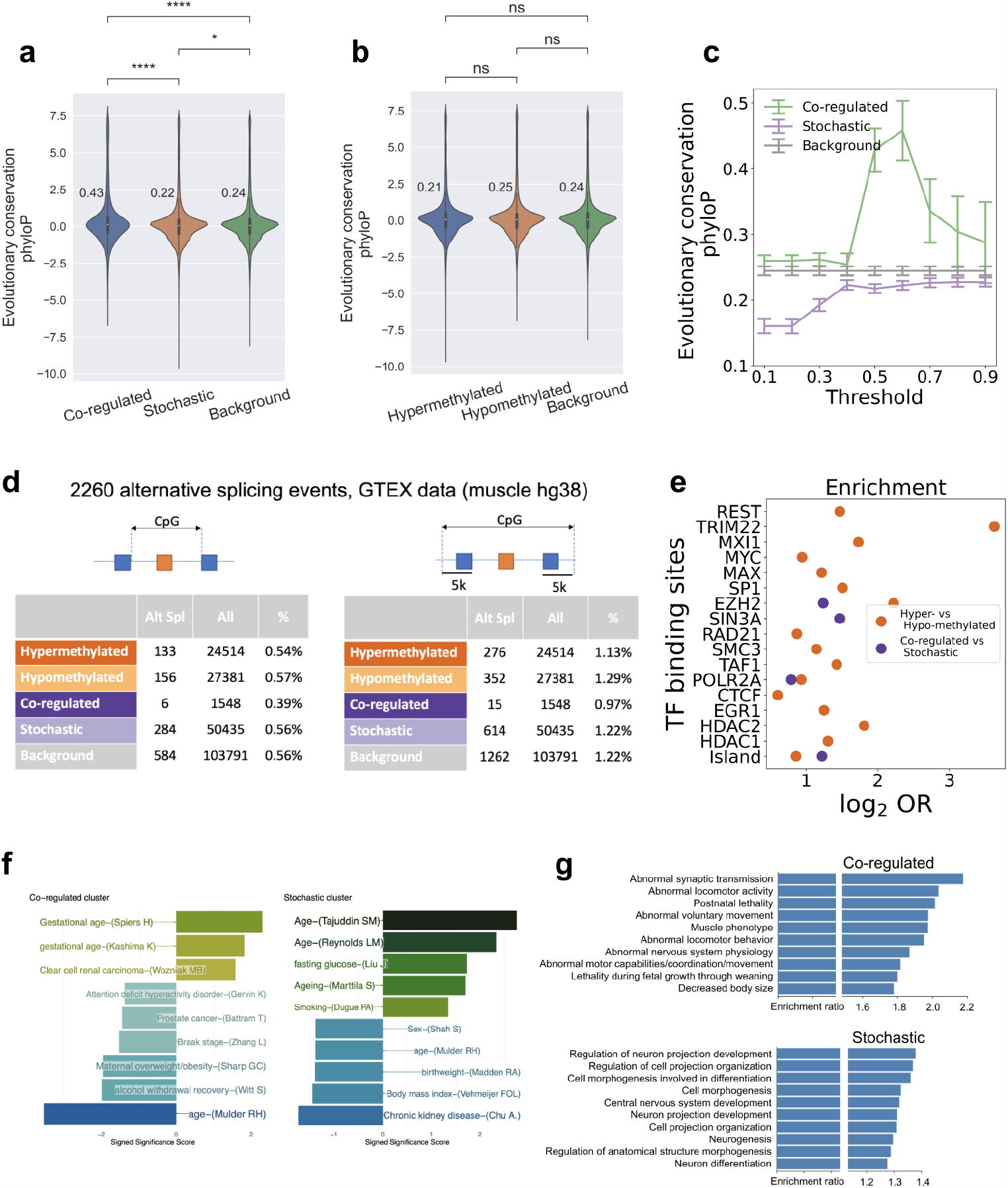
Biological annotation of co-regulated and stochastic clusters. **a**. PhyloP evolutionary conservation score distributions for CpGs comprising stochastic and co-regulated clusters compared to random regions of the genome. **b**. PhyloP evolutionary conservation score distributions for CpGs comprising hypermethylated and hypomethylated clusters compared to random regions of the genome. **c**. Evolutionary conservation score (phyloP) for co-regulated, stochastic and random sites as a function of the correlation threshold for 51,895 CpGs passing the coverage filter (5 cells of young mice, and 5 cells of old mice measured simultaneously). **d**. Enrichment analysis of age-associated alternative splicing events (Alt Spl) for the CpGs comprising co-regulated, stochastic, hypermethylated and hypomethylated clusters (**left**), and for alternative splicing events within a 5 kb distance of the CpGs clusters in the genome (**right**). The difference was not statistically significant (Fisher’s exact test) due to the scarcity of alternative splicing events. **e**. Enrichment with transcription-factor (TF) binding sites and CpG islands for co-regulated clusters vs. stochastic clusters and hypermethylated vs. hypomethylated regions. All shown TFs passed the level of statistical significance was chosen at 0. 05 after the Bonferroni correction for multiple comparisons. **f**. Enrichment of co-regulated and stochastic clusters against EWAS hits. Each horizontal bar represents an enriched term. The X-axis shows the -log10(P-value), signed by log2 (Odds ratio). Only the EWAS trait with significant enrichment (P < 0.05) are included and annotated. **g**. GO/KEGG enrichment analysis of genes associated with co-regulated and stochastic clusters: top-10 categories with the lowest false discovery rate (FDR < 0.05) across biological processes, cellular components, molecular function, KEGG pathways and mammalian phenotype ontology. *Mann-Whitney-Wilcoxon test p-values for legend in* **a** *and* **b**: *ns:* 0. 05 < *p* ≤ 1, *\*:* 0. 01 < *p* ≤ 0. 05, *\*\*:*10 ^−3^ < *p* ≤ 0. 01, *\*\*\*:* 10^−4^ < *p* ≤ 10^−3^, *\*\*\*\*:* . 0. 05 < *p* ≤ 10^−4^

We further examined age-associated splicing events in the muscle tissue from the Genotype-Tissue Expression (GTEx) database^57^. For each CpG site from the co-regulated, stochastic clusters and random regions, we checked if it was located in the region from the beginning of the first exon to the end site of the last exon in an alternative splicing event (**Fig. 6d left**), or surrounding the splicing event within 5 kb (**Fig. 6d right)**. Co-regulated clusters showed fewer alternative splicing events in comparison with stochastic clusters for correlation threshold 0.6 (**Ext. Data Fig. 5** for correlation thresholds 0.4 and 0.5); however, the difference was not statistically significant (Fisher’s exact test).

We also examined enrichment of co-regulated clusters vs stochastic clusters and hypermethylated vs hypomethylated clusters with transcription factor (TF) binding sites (**Fig. 6e**), and compared the trends with random genomic regions (**Ext. Data. Fig. 2e**). Co-regulated regions contained significantly more EZH2, SIN3A, POLR2A binding sites and CpG islands than stochastic clusters. Distal location of co-regulated regions from main gene sequences may explain the observation of a lower number of alternative splicing events in the co-regulated regions in **Fig. 6d**.

We analyzed the enrichment of CpG sites from co-regulated and stochastic clusters for correlation threshold 0.6 (**Ext. Data Fig. 6** for correlation thresholds 0.4 and 0.5) with hits from 6,993 epigenome-wide association studies (EWAS)^58^. Co-regulated clusters were enriched with phenotypes related to gestational age and clear cell renal carcinoma, and stochastic clusters with phenotypes related to age, fasting glucose, aging and smoking (**Fig. 6f**).

The lists of genes associated with co-regulated and stochastic clusters were further subjected to GO/KEGG enrichment analysis^59^ (**Fig. 6g**). Co-regulated genes were enriched for muscle-specific pathways, including abnormalities of development, such as abnormal muscle, locomotion phenotypes, fetal and postnatal lethality and decreased body size. Stochastic genes were enriched for cell morphogenesis and neuron differentiation categories. The enrichment for muscle-specific pathways among co-regulated genes, given that the original dataset was measured in muscle stem cells, may signify the programmatic nature of those genes preserving cell identity, whereas the stochastic genes enriched for neuron and differentiation pathways may be responsible for the adaptation to stochastic changes in the cell microenvironment.

### Decreased epigenetic co-regulation is associated with lower transcriptomic coordination in multiomics data

To test the hypothesis that epigenetic co-regulation affects transcriptomic coordination, we examined the single-cell RNAseq data corresponding to the scDNAm data analyzed above from the multi-omic aging muscle stem cells (muSC)^49^. The transcriptional coordination of a set of genes was measured with the help of the global coordination level (GCL)^29,30^. First, we mapped the list of CpGs comprising co-regulated or stochastic clusters to genes (**Fig. 7a**). We considered a CpG associated with a given gene if it was mapped within the gene’s promoter, 5’UTR, exon, intron, exon/intron and intron/exon boundaries and 3’UTR region according to the genome annotation (mm10)^60^ on one of two DNA strands.

**Fig. 7.**
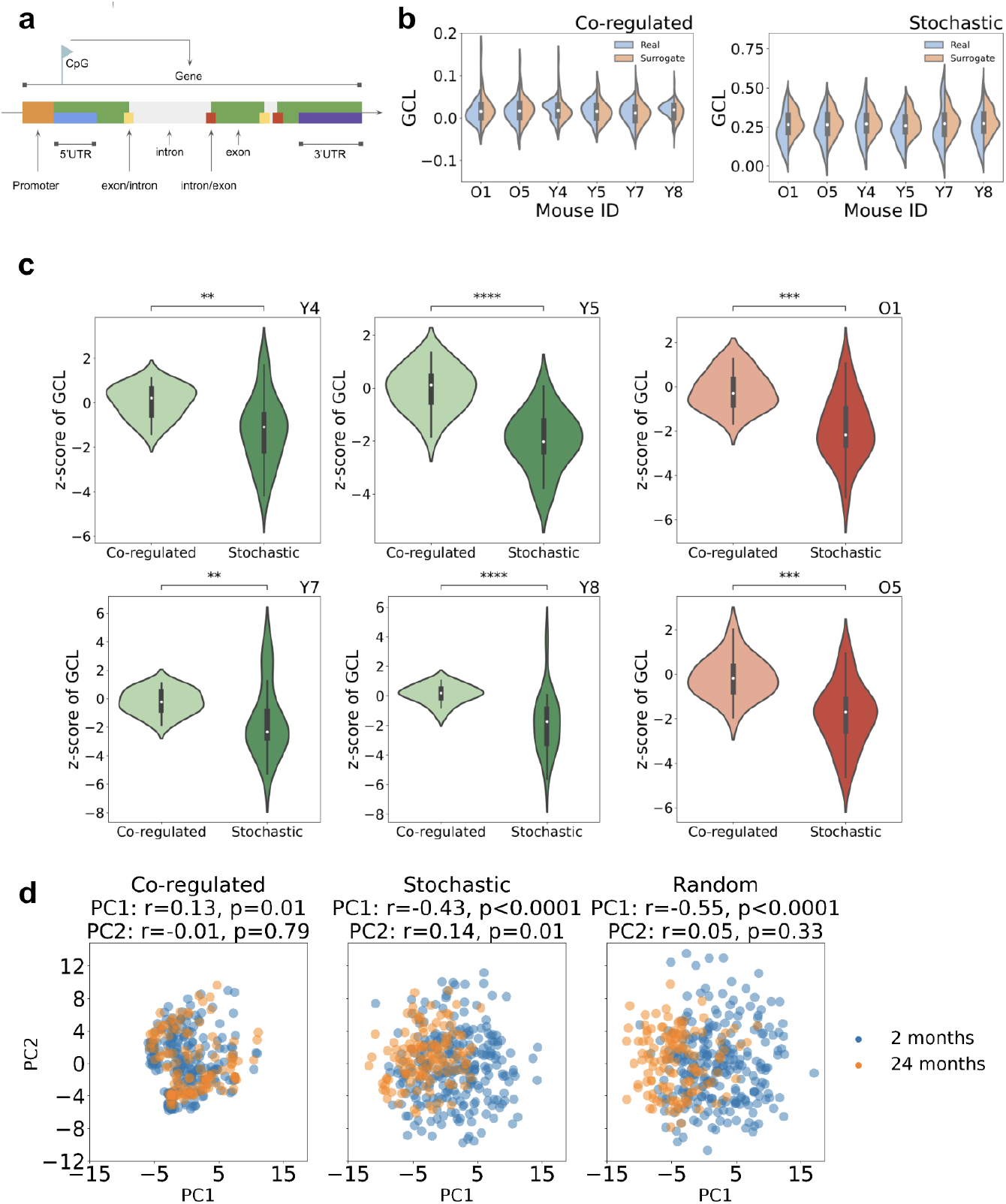
Decreased global coordination levels (GCL) for gene expression of genes associated with stochastic compared to co-regulated clusters. This analysis was conducted on the scRNAseq part of the multiomic dataset from **Fig. 4. a**. Mapping of CpG sites comprising co-regulated and stochastic clusters to gene lists. **b**. Global coordination level of gene expression for genes associated with co-regulated and stochastic clusters of CpGs in young mice (2 months old: Y4, Y5, Y7 and Y8) and old mice (24 months old: O1 and O5). Blue distributions (“Real” in legend) represent true distributions of the GCL for the given gene sets, whereas “Surrogate” distributions are for surrogate gene sets that were randomly selected from a subset of genes with similar expression levels as the gene-set genes. Each surrogate gene set preserved the size of the original gene set and mimicked its expression profile. **c**. For normalization of results presented in **b**, we use Z-scores of the GCL relative to the corresponding surrogate gene sets (see Methods for details). Co-regulated genes show a significantly higher Z-score than the stochastic ones for all mice. The results in **b** and **c** are shown for the sense DNA strand, see **Ext. Data. Fig 3** for the antisense DNA strand results. **d**. Principal component analysis for co-regulated, stochastic and random genes. The Pearson correlation coefficients of PC1 and PC2 with age are shown in the titles. The co-regulated genes have the smallest correlation with age implying a tighter control during aging and supporting their epigenetic co-regulation.

For the sense DNA strand, we calculated the GCL for co-regulated and stochastic gene lists for each mouse. We compared the GCL value of each geneset (co-regulated and stochastic) to the GCL values of their corresponding “surrogate” gene sets. These surrogate gene sets were randomly selected from a subset of genes with similar expression levels preserving the size of the original gene set and mimicking its expression profile (see Methods for more detail). By analyzing the distribution of GCL values of the gene set compared with the distribution of the surrogate gene sets, we can test whether the former tend to have elevated/lower coordination (**Fig. 7b**). In doing so, we calculated the Z-scores of GCL against the corresponding surrogate gene set: 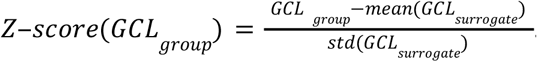 Comparison of GCL Z-scores showed that genes associated with co-regulated CpG clusters exhibit significantly higher Z-scores of GCL than those associated with stochastic CpG clusters: in both 2-month-old and 24-month-old (**Fig. 7c**) mice. The same dependence was observed for the other DNA strand (**Ext. Data Fig. 3**).

We also tested if co-regulation affected aging changes in scRNAseq signal (see Methods) by plotting the principal first and second components for co-regulated, stochastic and random subsets of genes against age (**Fig. 7d**). The co-regulated genes show the smallest correlation with age compared to the stochastic and random genes. This suggests that their tight control did not change during aging, and the gene expression profile remained similar, which supports their epigenetic co-regulation.

Overall, the GCL analysis of scRNAseq data is in a good agreement with the scDNAm clustering: in most cases, co-regulated CpG clusters are associated with the group of genes showing transcriptomic coordination similar to their surrogate gene sets, whereas the stochastic ones showed significantly lower GCL than their surrogate. Moreover, after normalization to the z-scores against their surrogate lists, the Z-scores of GCL for the co-regulated gene set were significantly higher than the Z-scores of the stochastic gene set for all mice. This analysis demonstrates that the loss of epigenetic co-regulation is associated with the loss of transcriptomic coordination. Our analysis, though limited by the currently available data, implies that the loss of regulation during aging universally spans at least two levels of regulation: epigenetic and transcriptional. However, investigation of causal relations between epigenetic co-regulation and transcriptomic coordination would require further research in different cell types and organisms.

## Discussion

We analyzed the dynamics of single-cell and tissue DNAm during functional aging and embryonic development, with the goal of uncovering and separating stochastic accumulation of DNAm changes from co-regulated DNAm changes driven by a common biological process. We also analyzed global coordination level (GCL)^29,30^ of genes based on single-cell RNAseq data matching the scDNAm aging dataset^49^. By examining tissue DNAm changes with age and the respective DNAm aging clocks, we found that epigenetic aging is an omnipresent (for example, 13.6% of measured CpGs changed significantly with age) yet a relatively slow process. It is also very small in amplitude (most sites change ≤ 10%), is characterized by common temporal dynamics and shows increased heterogeneity with age. By simulating a null-hypothesis that DNAm aging changes occur randomly in single cells, we managed to reproduce the experimental aging clock dynamics by simulation of a stochastic decay model. All these observations are consistent with the concept of epigenetic aging being a stochastic process characterized by increasing entropy.

In order to test whether other aging dynamics are also present in DNAm data, we turned to aging scDNAm and found that 92% of the measured CpGs behaved in a stochastic manner during aging, whereas 8% changed in a co-regulated way. An scRNAseq GCL analysis of genes associated with CpGs in co-regulated and stochastic clusters indicated that lower epigenetic co-regulation is associated with lower transcriptomic coordination, thus extending the concept of co-regulation beyond the scope of epigenetics to transcriptomics and supporting the hypothesis of the existence of a shared biological mechanism responsible for epigenetic co-regulation and transcriptomic coordination during aging (see Fang *et al*.^61^ for a study of the role of DNAm entropy during embryonic development). During gastrulation scDNAm changes were dominated by a global co-regulated methylation event, in the background of which, due to low sequencing quality and high sequencing noise, we managed to identify fewer than 5% of additional co-regulated CpGs. Even though the available scDNAm data are currently scant to provide more detailed insights into epigenetic aging, the methods developed in the present paper would be applicable to future single-cell data generated with advanced sequencing techniques. In particular, the algorithm we developed for the identification of co-regulated CpG clusters may allow improving the accuracy and interpretability of DNAm aging clocks. However, additional research would be required to clarify the mechanisms responsible for epigenetic co-regulation and transcriptomic coordination and how they might affect each other during aging. Moreover, the loss of regulation during aging should also have an effect on multiple organismal levels (DNA mutations, epigenetic drift, transcriptional noise, etc.), and it would be interesting to extensively examine this in future research.

We applied typical epigenetic clock-building routines (Lasso and ElasticNet penalized regressions) in order to build epigenetic aging clocks and showed that they may be biased towards stochastic CpG clusters. Without lowering accuracy of chronological age predictions, they may ignore most of the co-regulated CpG sites due to their high collinearity. In response to an intervention, co-regulation of a cluster may be disrupted, which may be missed by such clocks. It is likely that stochastic CpG sites bear less information regarding the biology involved because of their high tolerance to stochastic epigenetic changes. In other words, the clocks built to measure stochastic accumulation of epigenetic changes, might not perform well where one expects reversal of biological pathways and processes, for example, in the case of rejuvenation therapies. On the other hand, clocks built on the co-regulated cluster of CpG sites may perform worse in the context of chronological age prediction but may be able to better capture the effects of longevity interventions.

Both tissue and single-cell DNAm analyses suggest that the high accuracy of epigenetic aging clocks may be predetermined in part by the stochastic decay of the epigenetic state set during early development. We describe a mechanism for epigenetic clocks which is strikingly similar to radiocarbon decay often used for dating in archeology: there is no need to have a biologically relevant mechanism as soon as the mean concentration of radioactive carbon or of the fraction of methylated DNA change monotonically with age. Radiocarbon dating works surprisingly well even though it is based on a purely stochastic process of radioactive decay according to the exponential decay law. In contrast to the radioactive decay of carbon-14, in the case of epigenetic clocks, we deal with two dynamic processes of gaining methylation in some genomic regions and losing methylation in others^62^. Thus, the analogy with radiocarbon decay is rather mathematical and conceptual rather than biological. The two processes of loss and gain of methylation are controlled by two different kinds of biological machinery, but the measured mean methylation level changes are affected by both processes. Stochasticity implies that over time those machineries unavoidably make mistakes, which accumulate gradually with age and can be used as robust predictors of age. Additionally, organisms adapt to age-related changes through cell, tissue and organismal-level responses, again distinguishing them from inanimate objects. Adaptive changes may also be expected in response to stochastic damage.

It is important to note that our analyses are limited to the process of functional aging, and do not consider the effects of rejuvenation therapies on the epigenome^63–65^. Stochasticity of age-related epigenetic changes does not imply the impossibility of reversal, as is the case for epigenetic reprogramming protocols resetting the DNAm patterns. At the same time, stochasticity behind the process of accumulation of epigenetic changes with age does not preclude programmatic behavior, a quasi-program of aging, defined by the developmental biology predisposing species to follow a particular aging trajectory. Thus, components of the stochastic part of epigenetic clocks would be predetermined by development and biological organization of the organism. The sites that were initialized in the hypo- or hyper-methylated states during early development would tend to stochastically gain or lose methylation with age, hence they would make good candidates for epigenetic clock CpG sites. Thus, the developmental program initializes the epigenome into a state that later stochastically decays, under the direction of underlying biology, during aging. Therefore, the multispecies epigenetic clocks^66^ may work well because closely related species, such as mammals, share the associated developmental biology setting them into similar initial states of the epigenome. Overall, the effects of early embryonic development on aging need to be further investigated^40,67^.

The question of the biological meaning of existing epigenetic clocks deserves separate discussion. The causal relationship between molecular changes during aging and functional decline resulting in mortality is also the subject of an ongoing debate. A highly desirable feature of aging clocks is the ability to predict mortality events and lifespan; however, the state-of-the-art epigenetic clocks continue ticking in immortalized cell cultures^68,69^ and in naked-mole rats^70,71^, where mortality exhibits minimal changes with age. These observations raise questions about the use of existing epigenetic clocks, trained for chronological age, for mortality prediction. At the same time, the absence of a mechanistic explanation behind epigenetic clocks impedes their clinical use as aging biomarkers.

Stochasticity behind the clock does not imply that there is no biological value in the clock. Stochastic accumulation of damage is chiefly influenced by genetics and may be influenced by lifestyle, diet, interventions and other factors, hence it is possible both to reset stochastic changes to some other state (younger or older) and change the rate of damage accumulation (in both directions, up and down). Therefore, stochasticity of epigenetic clocks might be a good indicator of cumulative damage, with the contribution of cumulative deleteriousness of the environment in which an organism lives, or of cumulative non-specific damage. However, it is less clear how stochastic epigenetic clocks would capture the effects of target-specific therapies or some non-promiscuous aging changes. We anticipate that there might be two different kinds of clocks necessary for the quantification of aging: stochastic for the estimation of cumulative damage, and co-regulated for the estimation of programmatic effects of longevity interventions. Current approaches for building epigenetic clocks mix up these two qualitatively different components, limiting their predictive power for testing interventions.

### Limitations of the study

Apart from a technical limitation of our study due to the high sparsity of scDNAm datasets produced by the state-of-the-art sequencing methods, we note other non-technical limitations below. Any technical variation and noise in the data would count against our definition of co-regulation, however, we extensively tested the stability of our algorithm (see Methods, **Ext. Data Fig 8**) and showed that it is stable to a large proportion of missed measured and/or incorrectly labeled CpGs. More complex patterns of co-regulation, such as negative regulation, time-delayed feedback motives, allelic expression, noise regulons, and others, may be missed by the definition of co-regulation used in the present study due to the lack of high quality scDNAm data. Also, the fact that a random stochastic model can reproduce empirical distributions of tissue DNAm is not sufficient to state that the generative process is random and stochastic, yet we use it here as the first-order approximation of reality (see, for example, a transcriptomic change initially interpreted as “noise”, later proved to be deterministic^72^). Deterministic single-cell heterogeneity may account for a part of stochastic changes identified herein. There may be inherent sources of noise in scDNAm levels similar to those observed in transcriptional signals^73^, which we cannot account for with the currently available data. To completely rule out the possibility of mis-classification of some co-regulated CpG sites as stochastic, one would need to have access to a technique of longitudinal *in vivo* targeted non-destructive measurement of DNAm levels in single cells, which is yet to be developed. Currently available scDNAm techniques, such as bisulfite sequencing used in the present study for conceptual and illustrative single-cell analysis, are not yet capable of producing datasets of sufficiently high coverage and quality for a complete resolution of epigenetic aging dynamics. We leave this for future research when more advanced scDNAm sequencing technologies become available.

## Acknowledgements

The authors thank Didac Santesmasses, Jeyoung Bang, Wayne Mitchell, Anastasia Shindyapina, Alex Trapp for discussion. The work was supported by NIA grants, Impetus grant program and James Fickel and Michael Antonov Foundations.

## Code availability

The code used to produce the results and figures in the study is published on Github https://github.com/TarkhovAndrei/scDNAm.

## Supplementary Information

### Methods

#### Single-cell stochastic model of DNAm changes

For each CpG site, we assume the aging trajectory of its methylation level is controlled by three parameters: *q* — the initial methylation level, *p* _*d*_ — the rate of demethylation in a unit of time, *p*_*m*_ — the rate of methylation in a unit of time. The model is assumed to be purely stochastic, which means that the state of a CpG site in a short time interval probability to get methylated *p*_*m*_ ∆*t* and probability to get demethylated *p* _*d*_ ∆*t*.

Assuming that there are *n* cells in a sample, we can denote by *n*_*d*_ and *n*_*m*_ the numbers of cells that are demethylated or methylated for a given CpG site. The aging dynamics for *n*_*m*_ and *n*_*d*_ would be describable by the following rate equations:

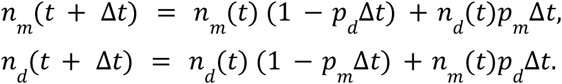

Conservation of the total number of sites is satisfied, which is shown by the summation of the two equations above: *n*_*m*_ (*t* + ∆*t*) + *n*_*d*_ (*t* + ∆*t*) = *n*_*m*_ (*t*)+ *n*_*d*_ (*t*) = *n*. To derive the differential equation for the average methylation level 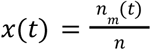, we use the fact that *n*_*d*_ (*t*) = *n* − *n*_*m*_ (*t*),

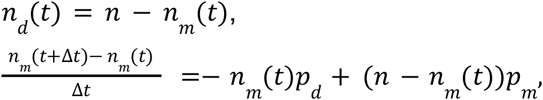

and by dividing both sides of the equation by *n*, we obtain:

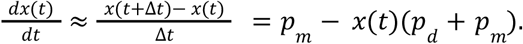

The exact solution of the above equation reads

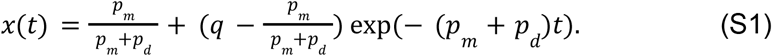

To understand the meaning of the three introduced parameters *q, p*_*m*_, *p* _*d*_, let us further analyze the above equation. The initial value of the average methylation level is defined by *x*(0) = *q*, whereas with time the methylation level tends to the asymptotic value 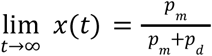. The rate of exponential decay is equal to the sum of the rates of methylation and demethylation *p*_*m*_ + *p*_*d*_.

It is worth noting that even though the process generating the dynamics is purely stochastic, it doesn’t necessarily lead to the saturation of the methylation level at the level 0.5. To the contrary, by varying the three parameters we may obtain an aging trajectory of a CpG site starting at any point from 0 to 1, and tending to any other methylation level from 0 to 1, whereas the rate of change would be controlled by the absolute values of the methylation and demethylation rates for each particular site. The above analysis considers a single CpG site and its aging trajectories. The values of the model parameters *q, p*_*m*_, *p* _*d*_ may be characteristic of a genomic position, and may bear some biological meaning.

#### Seven clusters of tissue DNAm changes during aging

In **Fig. 1g**, we clustered typical trajectories of DNAm level changes during aging into seven broad categories based on their initial methylation level and the direction of changes: three clusters not changing significantly (no significant Pearson correlation between DNAm levels and age, *p* > 0. 05) with age by their initial DNAm level for dynamics 1, 4 and 7 in the intervals [0; 0. 25], [0. 25; 0. 75] and [0. 75; 1], respectively. Dynamics 2 and 5 trend towards the methylation level of 0.5, while starting in the intervals [0; 0. 25] and [0. 75; 1],respectively. Dynamics 3 and 6 start in the interval [0. 25; 0. 75] and trend up and down, respectively. For dynamics 2, 3, 5 and 6, DNAm levels had a significant Pearson correlation with age, *p* < 0. 05.

#### Fitting experimental tissue DNAm aging trajectories to stochastic model prediction

In order to fit the experimental aging trajectories for CpG sites comprising the Petkovich *et al*. clock, we use the three-parametric stochastic aging trajectory derived above in Eq. (S1), and apply Python’s fitting tool scipy.opti*m*ize.curve_fit^74^.

#### Simulating random subset of aging trajectories predicted by the stochastic model

To show how a subset of aging trajectories corresponding to randomly sampled parameters *q, p*_*m*_, *p* _*d*_, we use Eq. (S1) and randomly draw the parameters from the uniform distributions defined on the following intervals: *q* ∈ [0, 1], *p*_*m*_, *p* _*d*_ ∈ [0, 0. 0015]. The number of sampled sets is equal to the number of CpG sites in the Petkovich *et al*. clock.

The bundle of random trajectories is thus fully defined by a single parameter — the upper bound for the rates *p*_*m*_, *p* _*d*_, which is set here to 0. 0015. The maximal rate of exponential decay among all CpG sites (*p*_*m*_ + *p* _*d*_) _*max*_ hence represents the critical parameter of the stochastic model. It might be related to the typical level of deleteriousness of the environment for the organism. Therefore, (*p*_*m*_ + *p*_*d*_)_*max*_ might be a proxy to the identification of the maximal lifespan for a species.

#### The role of sequencing coverage depth on tissue DNAm changes

To study the role of sequence coverage depth on the aging changes of DNAm levels, we repeated the analysis for **Fig. 1c**, for CpG sites with higher and lower coverage. Originally, we used 10X coverage depth, and tested whether 30X coverage significantly changes the maximum change of DNAm aging changes (**Ext. Data Fig 7)**. We used those CpGs that have 30X coverage in more than 200 mice out of 255 mice (high coverage limit), and fewer than 50 mice out of 255 mice (low coverage limit). The analysis indicates that lower coverage leads to higher amplitudes of DNAm changes thoughout lifespan, but we did not find any qualitative difference between 10X and 30X (high coverage) regimes.

#### Cell-type deconvolution for tissue DNAm signals

To test whether our predictions may be affected by cell-type composition changes, we used the web-interface for EpiDISH^75–77^. In order to do so, we converted mouse genome positions (mm10) to human positions (hg38) with the help of the UCSC Genome Browser liftover tool^78^, and used the closest available probe for the EPIC chip for each CpG. A converted table of DNAm levels for CpG sites was uploaded to EpiDISH for cell-type deconvolution. The transformation of mouse genomic positions to the human EPIC probes is not a one-to-one lossless procedure, and can potentially distort the cell-type deconvolution. Moreover, EpiDISH was designed to be used for human samples. We assume that the results produced by EpiDISH would be, at least partially, translatable to our mouse blood samples.

We normalized the fractions of cell types by the maximum value throughout the mouse lifespan. Our analysis of cell-type deconvolution for the CpGs whose DNAm levels changed significantly after the Bonferroni correction for multiple comparisons (**Ext. Data Fig 7c** and **d**) did not show any significant Pearson correlation for B, CD4T, CD8T cells, but showed a significant negative Pearson correlation for NK (*r* =− 0. 28, *p* < 0. 001), neutrophils (*r* =− 0. 60, *p* < 0. 001), monocytes (*r* =− 0. 58, *p* < 0. 001) and eosinophils (*r* =− 0. 60, *p* < 0. 001). This observation is consistent with previous observations for cell composition of PBMCs in C57BL/6J mice^79^. The typical relative change of neutrophils, monocytes and eosinophils is on the order of 30%, which is similar to the largest relative change of DNAm levels (**Fig. 1c, Ext. Data Fig 7a** and **b**). The similarity may indicate that the strong changes causing the nonlinearity observed in the DNAm dynamics can be accounted for by the cell-composition changes. However, given that the cell-decomposition analysis directly used the same DNAm information as was used in the prior analysis of the aging dynamics of DNAm levels, the observation and reasoning it causes may be circular. To exhaustively account for cell-composition changes, one should have access to cell sorting data associated with DNAm sequencing which was not performed in the original paper^4^.

#### Genomic enrichment analysis for epigenetic profiles

For genomic annotation of epigenetic profiles we used R package annotatr^60^.

#### Single-cell DNAm data analysis

Due to the high sparsity of single-cell DNAm data, we had to extensively use filtering by coverage. For illustrative purposes of identifying visually recognizable co-regulated clusters, we set a threshold of coverage for each CpG for it to be measured in 15 young cells and 15 old cells. In each of the cells, we had to keep all CpGs covered by at least 1 NGS read due to low coverage. For enrichment analysis, we lowered the filter to 5 young and 5 old cells. For muSCs, one old mouse had to be censored because of an extremely low coverage. For embryonic scDNAm, we set the threshold at 25 covered cells for each embryonic stage for each CpG.

For correlation analysis we developed custom correlative algorithms that were able to ignore omitted CpGs in some cells, and use only those that were measured. To avoid the confounding factor of different coverage in young and old cells for identifying CpGs whose methylation significantly correlated with age, we also produced a pseudo-bulk sample for each of the mice by calculating the average methylation level for each of the CpGs. A CpG was considered significantly correlated with age if it was associated with age in both single cell data, and in the pseudo-bulked data.

For inter-cell Pearson correlative analysis, we used a custom Python code taking into account only those methylation values that had a non-zero coverage. Non-measured CpGs with zero coverage were not used for calculating the Pearson correlation. The threshold correlation value for a pair of CpGs to be considered co-regulated was chosen to be in the range from 0.1 to 0.9, which must be satisfied in three groups of mice: only young mice, only old mice and all mice. For embryonic data, there were four groups corresponding to each of the embryonic days E4.5-7.5. That further allowed lowering the limit for false positive identification of co-regulation in the case of low coverage.

#### Algorithm for identification of co-regulation in sparse scDNAm data

For identification of co-regulated clusters in sparse scDNAm data, we developed an algorithm based on the inter-cellular Pearson correlation for pairs of CpG sites. The algorithm takes into account a high level of sparsity due to the low coverage of sequencing.

For practical calculations, we use the Pearson correlation coefficient of a pair of CpGs defined as 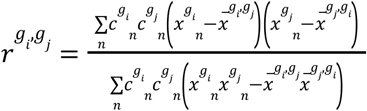, where the unbiased mean methylation level for CpG *g*_*i*_ over pairs of cells where CpGs *g*_*i*_ and *g*_*j*_ were measured simultaneously was defined as 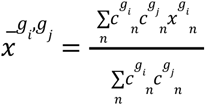 . for *i* = 1, 2. The sparsity mask defined as 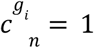 if CpG *g*_*i*_ in cell *n* was covered by a read, otherwise 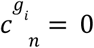.

A pair of CpGs was called “co-regulated” if the inter-cell Pearson correlation was larger than a given threshold correlation value in the range from 0.1 to 0.9, which had to be satisfied in three groups of mice (only young mice, only old mice and all mice) for the aging studies, and at each embryonic stage and all stages combined for the embryonic studies.

We extensively tested this algorithm on simulated data (**Ext. Data Fig. 8**). The simulation assumed 40 young and 40 old cells, each having 200 CpGs forming 4 blocks of 50 CpGs. The first half of CpGs are stochastic, and the second half is co-regulated. In each group of 100 CpGs, 50 CpGs originally have the DNAm level of 30%, and reaching 70% in older cells, another 50 CpGs start at 70% and decrease methylation to 30%. We tested the algorithm’s stability to errors in DNAm sequencing (identifying sites methylated instead of demethylated, and *vice versa*), and missing values (having no coverage). The error rate was varied from 0 to 40%, and the missing values percentage from 0 to 90%. The algorithm exhibits robust identification for co-regulated CpGs up to 30% of errors, and 60% of missed values. The worst case scenario the algorithm handled well was 40% of missed values and 20% of errors. The algorithm tends to penalize co-regulation if the percentage of errors grows, whereas it favors co-regulation if we decrease the number of missed values. At the same time, the algorithm is stable to changes of the threshold of correlation, when 10% of errors are present in a wide range of percentages of missing values (up to 60%). Overall, the simulation check confirmed that the algorithm has met the necessary criteria.

#### Enrichment analysis in aging-associated splicing events

To detect age-associated splicing events, we downloaded RNA-sequencing files of muscle tissue from Genotype-Tissue Expression (GTEx) database^57^, quantified all alternative splicing events and modeled the association with age using linear regression. In total, 2,260 aging-associated alternative splicing events were identified. For each CpG site, we checked if the CpG site fell in at least one aging-associated splicing event (from the start site of the first exon to the end site of the last exon in an alternative splicing event), or surrounding the splicing event within 5kb. We counted this type of CpG site in each CpG cluster and compared it with random regions of the background genome.

#### Transcription factor binding regions

We used the ENCODE transcription factor binding database^80^ for identification of genomic regions corresponding to transcription factor binding sites. The database is based on ChIP-seq combining chromatin immunoprecipitation with DNA sequencing to infer possible binding sites of DNA-associated proteins. Prior to use, mouse CpG sites from the mm10 reference mouse genome were mapped to the conserved human CpG sites in the hg38 human reference genome with the help of the UCSC Genome Browser liftover tool^78^.

#### Enrichment analysis in phenome-wide EWAS signals

We analyzed the enrichment of CpG sites from co-regulated and stochastic clusters in hits from 6,993 epigenome-wide association studies in humans, obtained from the EWAS catalog^58^. Mouse CpG sites from the mm10 reference mouse genome were mapped to the conserved human CpG sites in the hg19 human reference genome with the help of the UCSC Genome Browser liftover tool^78^. Then, the nearest human CpG sites in the Illumina EPIC array within 100 base-pairs were identified and used for the enrichment. For each EWAS hit, Fisher’s exact test was performed to determine the enrichment of either co-regulated or stochastic clusters for a given trait. We did not correct our EWAS analysis for potential biases due to the selectivity of RRBS to CpG dense regions, and the DNAm arrays to CpG inclusion selection^72^.

### Global Coordination Level (GCL)

Consider an *N* × *M* matrix representing the gene expression of *N* genes from *M* single cells, i.e., a matrix element *X*_*i,j*_ represents the expression level of gene *i* in cell *j*. To quantify the global coordination level, we first divide the genes into two random subsets, *S* ^*k*^_1_and *S* ^*k*^_2_ each of them consists of *N*/2 genes. Then, we measure the dependency level *D k* = *D*_*bcDcorr*_ (*X*_1_, *X*_2_) where 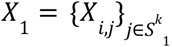 and 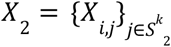 . In this study, we use *bcdCorr*, the dependency level between two high-dimensional variables, described in detail before^29^, but essentially any high dimensional dependency measure can be used (e.g., mutual information). We repeat these steps *m* times and define the GCL as the average of the dependency levels 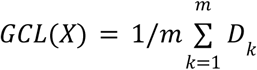

Previously^29^, we showed that the GCL stabilizes for m>50. Accordingly, in our analysis we choose m=50.

### Single-cell RNAseq data analysis

In single-cell RNAseq data (scRNAseq), we analyzed GCL of three different clusters in five cohorts of cells obtained from six different mice. Gene expression is represented as the logarithm of normalized counts, i.e., transcript per million (TPM), for each cell, *X*_*i,j*_ = *log*_2_ (*TPM* _*i,j*_ + 1). We applied the following routine across all cell cohorts: (a) Remove genes that are expressed (*X*_*i,j*_ >2) in fewer than 10% of the cells in each cohort. (b) Filter cells for which the number of expressed genes (*X*_*i,j*_ >2) is larger or smaller than two standard deviations from the mean number of expressed genes. We analyzed two sets of genes: “Stochastic” and “Co-regulated” that were mapped from scDNAm data according to the mapping described in the main text. Each of these sets had a different size and a different gene expression profile, with different mean expression values. In order to reduce possible biases due to the size and average gene expression profile, when comparing the GCL of these groups, we have sampled each group in a way that will ensure similar gene set sizes and mean gene expression profiles.

We normalized the GCL value of each geneset to the GCL values of corresponding ‘surrogate’ gene sets (“Co-regulated” and “Stochastic”). These surrogate gene sets were randomly selected from a subset of genes with similar expression levels as the gene-set genes. Thus, each surrogate gene set preserved the size of the original gene set and mimicked its expression profile (Extended Data Fig. 4d–f in Ref.^29^ for a detailed explanation of the surrogate procedure). By analyzing the distribution of GCL values of the gene set compared with the distribution of the surrogate gene sets, we can test whether the former tend to have elevated/lower coordination.

The normalized GCL score was calculated as a Z-score against the surrogate GCL distribution:

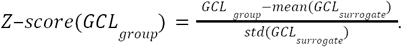

To produce **Fig. 7d**, we used co-regulated, stochastic and random gene lists to produce three subsets of our scRNAseq data. Then, we applied the log-normalization of counts in each dataset, and ran principal component analysis of the resulting data matrices. For plotting, we chose two first principal components, and calculated their Pearson correlation with age and its p-value.

## Extended Data

**Extended Data Fig. 1.**
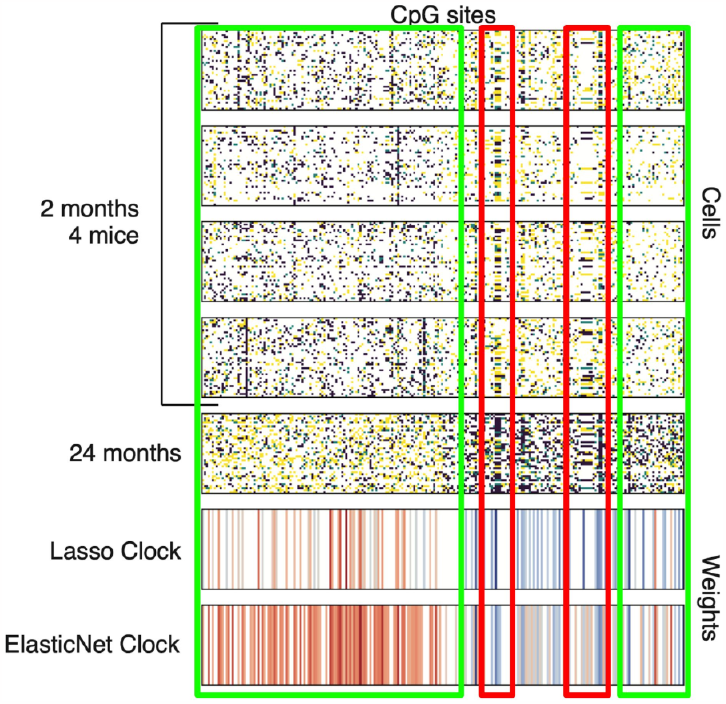
Dynamics of CpG sites significantly associated with age in scDNAm data^47^. The dominant cluster of 92% of all CpG sites changes stochastically (green boxes), whereas co-regulated clusters represent 8% of all CpG sites changing with age (red boxes). Penalized regression models are biased towards stochastic CpG sites, since they penalize any kind of correlation across the CpG sites used in the model (Lasso and ElasticNet clocks). Color represents the value of the regression weights. Co-regulated clusters contain fewer non-zero weights in the regression models. Therefore, the penalized regression models lower the fraction of co-regulated sites used for building DNAm aging clocks.

**Extended Data Fig. 2.**
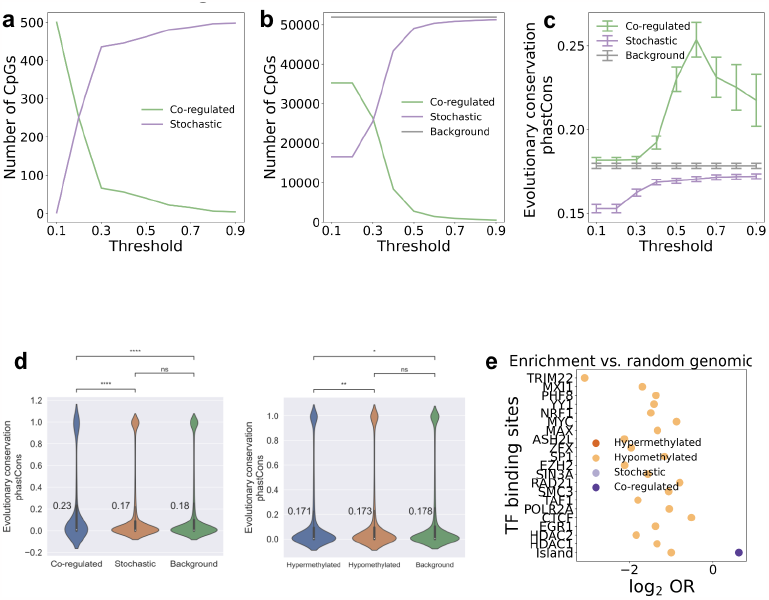
Biological annotation of co-regulated and stochastic clusters. **a**. Number of co-regulated and stochastic sites as a function of the correlation threshold for 502 CpGs passing the coverage filter (15 cells of young mice, and 15 cells of old mice measured simultaneously). **b**. Number of co-regulated and stochastic sites as a function of the correlation threshold for 51,895 CpGs passing the coverage filter (5 cells of young mice, and 5 cells of old mice measured simultaneously). **c**. Evolutionary conservation score (phastCons) for co-regulated, stochastic and random sites as a function of the correlation threshold for 51,895 CpGs passing the coverage filter (5 cells of young mice, and 5 cells of old mice measured simultaneously). **d**. phastCons evolutionary conservation score distributions for CpGs comprising stochastic and co-regulated clusters compared to random regions of the genome (**left panel**). Co-regulated clusters show significantly higher evolutionary conservation than the random regions, whereas stochastic clusters are significantly less conserved than both the random regions and co-regulated clusters. phastCons evolutionary conservation score distributions for CpGs comprising hypermethylated and hypomethylated clusters compared to random regions of the genome (**right panel**). Hypermethylated clusters show significantly lower evolutionary conservation scores than both the random regions and hypomethylated regions. **e**. Enrichment with transcription-factor (TF) binding sites and CpG islands for co-regulated and stochastic clusters, hypermethylated and hypomethylated regions vs. random regions of the genome. The level of statistical significance was chosen at 0. 05 after the Bonferroni correction for multiple comparisons.

**Extended Data Fig. 3.**
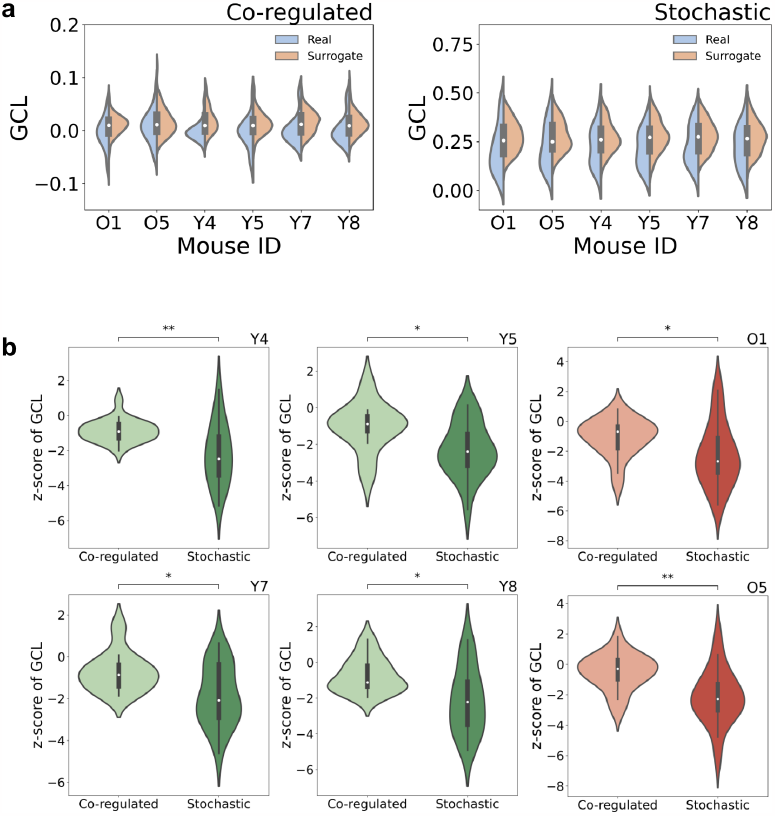
Global coordination levels for the antisense DNA strand. **a**. Global coordination level of gene expression for genes associated with the co-regulated and stochastic clusters of CpGs in young mice (2 months old: Y4, Y5, Y7 and Y8) and old mice (24 months old: O1 and O5). Blue distributions (“Real” in legend) represent true distributions of the GCL for the given gene sets, whereas “Surrogate” distributions are for surrogate gene sets that were randomly selected from a subset of genes with similar expression levels as the gene-set genes. Each surrogate gene set preserved the size of the original gene set and mimicked its expression profile, but did not represent any known KEGG pathway. **b**. For normalization of results presented in **a**, we use Z-scores of the GCL relative to the corresponding surrogate gene sets. Co-regulated genes show a significantly higher level of transcriptomic coordination than the stochastic ones for all mice. The results in **a** and **b** are shown for the antisense DNA strand.

**Extended Data Fig. 4.**
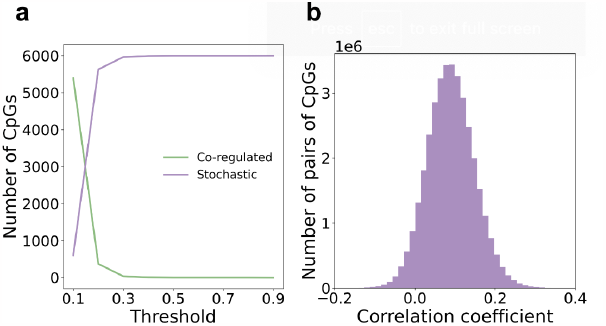
Co-regulation routine details for analyzing embryonic development. **a**. Number of CpGs in co-regulated and stochastic clusters for embryonic data as a function of the correlation threshold. **b**. Histogram of Pearson correlation coefficients for pairs of CpGs for embryonic development. The typical correlation coefficient is low, thus implying a high level of sequencing noise in the data.

**Extended Data Fig. 5.**
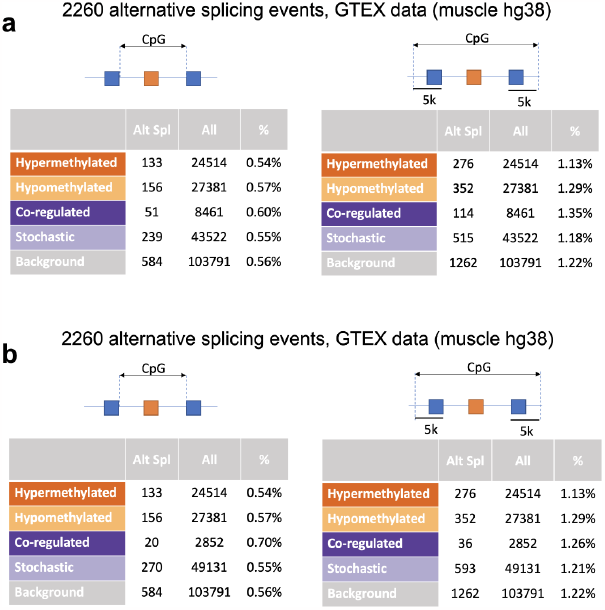
Enrichment for alternative splicing events. **a**. Enrichment analysis of age-associated alternative splicing events (Alt Spl) for the CpGs comprising co-regulated, stochastic, hypermethylated and hypomethylated clusters (**left**), and for the alternative splicing events within a 5 kb distance of the CpGs clusters in the genome (**right**). Co-regulated clusters showed a lower percentage of alternative splicing events than stochastic clusters, though the difference was not statistically significant (Fisher’s exact test). The correlation threshold is 0.4. **b**. Same as **a** for the correlation threshold 0.5.

**Extended Data Fig. 6.**
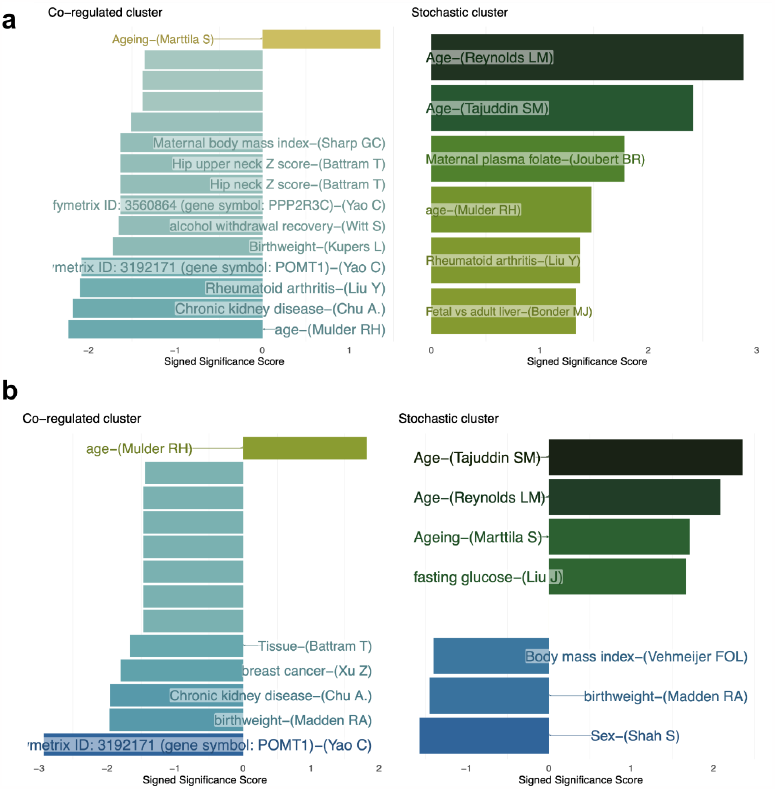
Enrichment of co-regulated and stochastic clusters against EWAS hits. Each horizontal bar represents an enriched term. The X-axis shows the -log10(P-value), signed by log2 (Odds ratio). Only the EWAS trait with significant enrichment (P < 0.05) are included and annotated. The results presented for correlation threshold: **a**. 0.4. **b**. 0.5.

**Extended Data Fig. 7.**
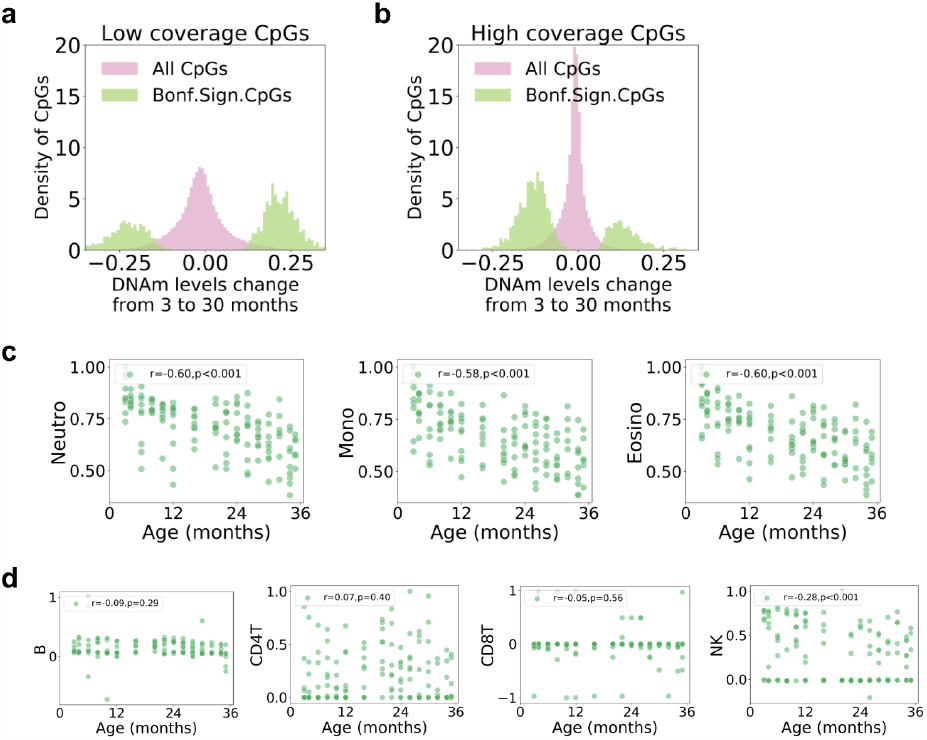
Role of coverage depth on DNAm changes during aging. The range of DNAm changes at CpGs sites in the dataset. Histograms are shown for all CpG sites and for the CpGs significantly changing with age. **a**. For CpG sites covered at the 30X coverage depth in no more than 50 mice out of 255 mice. **b**. For CpG sites covered at the 30X coverage depth in more than 200 mice out of 255 mice. **c**. Cell-type decomposition (EpiDISH) of tissue DNAm based on the CpG sites significantly changing with age after the Bonferroni correction for multiple testing. Neutrophils, monocytes and eosinophils were inferred to be decreasing significantly with age. All values were normalized to the maximum for a corresponding cell type. **D**. Same as **c** for B, CD4T, CD8T and NK cells. Only NK cells changed significantly with age.

**Extended Data Fig. 8.**
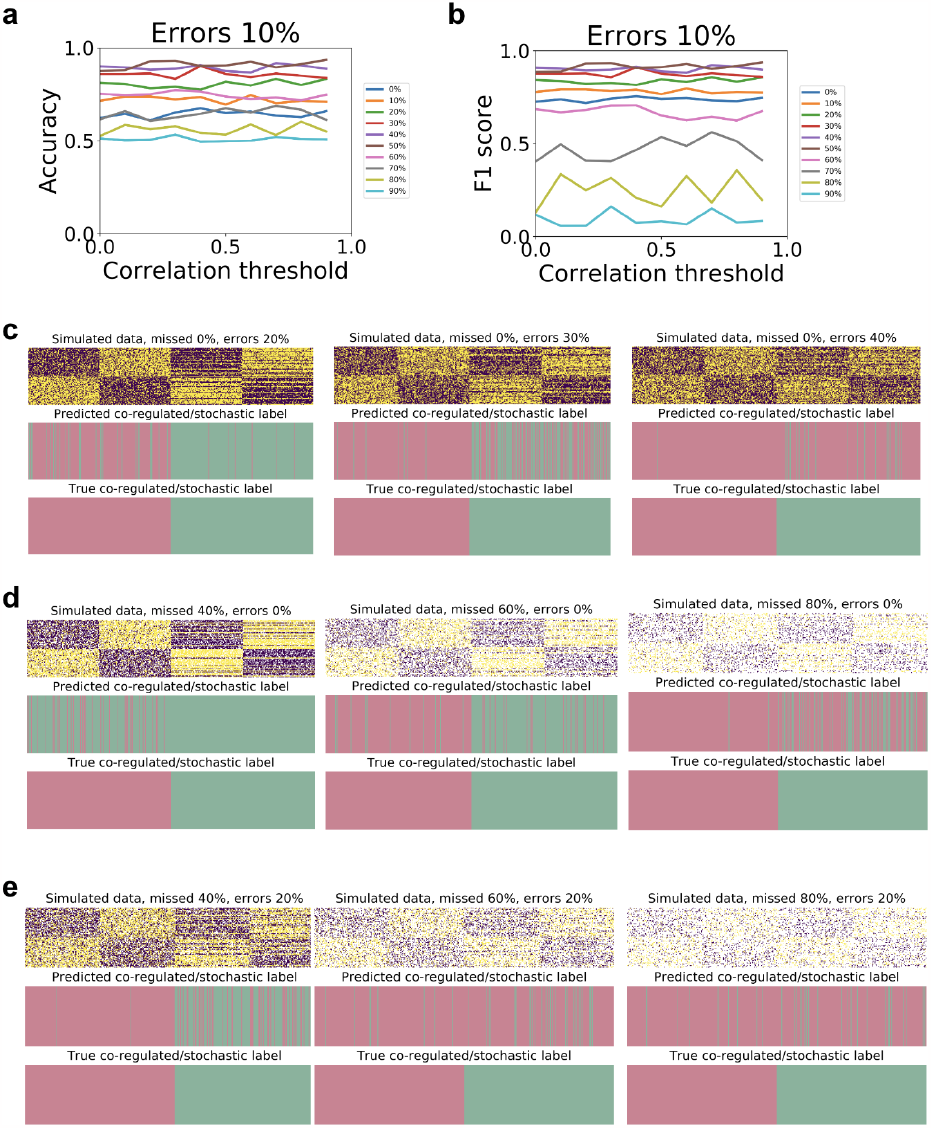
Tests of the algorithm for identification of co-regulated CpGs on simulated data. Simulation assumes 40 old and 40 young cells, each having 200 CpGs forming 4 blocks of 50 CpGs. The first half of CpGs are stochastic, the second half is co-regulated. In each group of 100 CpGs, 50 CpGs originally have the DNAm level of 30%, and reaching 70% in older cells, another 50 CpGs start at 70% and decrease methylation to 30%. The percentage of missed CpGs and the percentage of errors in identification DNAm level are varied, as well as the correlation threshold. **a**. Accuracy and **b**. F1 score of identification of co-regulated sites as a function of correlation threshold, assuming 10% of errors in DNAm levels, and missing values varying from 0% to 90% (in legend). **c**. Simulated data for 0% missed values, and errors varied from 20%, 30% and 40% respectively, along with the predicted co-regulated/stochastic CpGs (red for stochastic, green for co-regulated). **d**. Same as **c** for 0% of errors, and 40%, 60% and 80% of missed values. **e**. Same as **c** for 20% of errors, and 40%, 60% and 80% of missed values.

**Extended Data Fig. 9.**
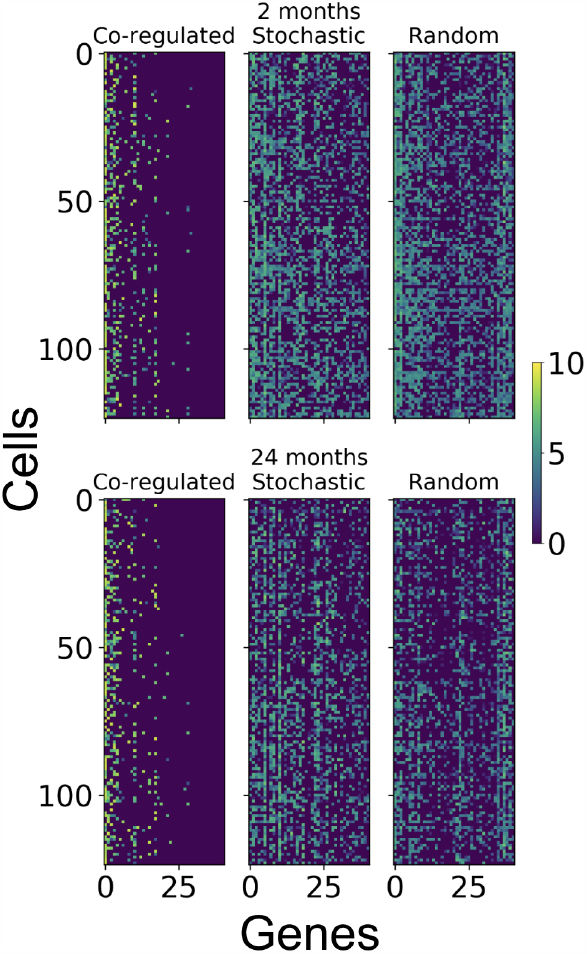
Heatmap of gene expression profiles for co-regulated, stochastic and random gene lists for young and old cells. Gene expression heatmaps are plotted for the top-41 genes ranked according to the first principal component weights from Fig. 7d. The gene expression levels were log-normalized.

## References

1. Horvath, S. DNA methylation age of human tissues and cell types. Genome Biol. 14, R115 (2013).

2. Hannum, G. et al. Genome-wide Methylation Profiles Reveal Quantitative Views of Human Aging Rates. Mol. Cell 49, 359–367 (2013).

3. Johansson, Å., Enroth, S. & Gyllensten, U. Continuous Aging of the Human DNA Methylome Throughout the Human Lifespan. PLoS ONE 8, e67378 (2013).

4. Petkovich, D. A. et al. Using DNA Methylation Profiling to Evaluate Biological Age and Longevity Interventions. Cell Metab. 25, 954-960.e6 (2017).

5. Thompson, M. J. et al. A multi-tissue full lifespan epigenetic clock for mice. Aging 10, 2832–2854 (2018).

6. Meer, M. V., Podolskiy, D. I., Tyshkovskiy, A. & Gladyshev, V. N. A whole lifespan mouse multi-tissue DNA methylation clock. eLife 7, e40675 (2018).

7. Levine, M. E. et al. An epigenetic biomarker of aging for lifespan and healthspan. Aging 10, 573–591 (2018).

8. Lu, A. T. et al. DNA methylation GrimAge strongly predicts lifespan and healthspan. Aging 11, 303–327 (2019).

9. Belsky, D. W. et al. DunedinPACE, a DNA methylation biomarker of the pace of aging. eLife 11, e73420 (2022).

10. Vershinina, O., Bacalini, M. G., Zaikin, A., Franceschi, C. & Ivanchenko, M. Disentangling age-dependent DNA methylation: deterministic, stochastic, and nonlinear. Sci. Rep. 11, 9201 (2021).

11. Seale, K., Horvath, S., Teschendorff, A., Eynon, N. & Voisin, S. Making sense of the ageing methylome. Nat. Rev. Genet. (2022) doi:10.1038/s41576-022-00477-6.

12. Tarkhov, A. E., Denisov, K. A. & Fedichev, P. O. Aging clocks, entropy, and the limits of age-reversal. bioRxiv (2022) doi:10.1101/2022.02.06.479300.

13. Levine, M. E., Higgins-Chen, A., Thrush, K., Minteer, C. & Niimi, P. Clock Work: Deconstructing the Epigenetic Clock Signals in Aging, Disease, and Reprogramming. http://biorxiv.org/lookup/doi/10.1101/2022.02.13.480245 (2022) doi:10.1101/2022.02.13.480245.

14. Bell, C. G. et al. DNA methylation aging clocks: challenges and recommendations. Genome Biol. 20, 249 (2019).

15. Jaffe, A. E. & Irizarry, R. A. Accounting for cellular heterogeneity is critical in epigenome-wide association studies. Genome Biol. 15, R31 (2014).

16. Marioni, R. E. et al. DNA methylation age of blood predicts all-cause mortality in later life. Genome Biol. 16, 25 (2015).

17. Tomusiak, A. et al. Development of a novel epigenetic clock resistant to changes in immune cell composition. http://biorxiv.org/lookup/doi/10.1101/2023.03.01.530561 (2023) doi:10.1101/2023.03.01.530561.

18. Kim, J. Y., Tavaré, S. & Shibata, D. Counting human somatic cell replications: Methylation mirrors endometrial stem cell divisions. Proc. Natl. Acad. Sci. 102, 17739–17744 (2005).

19. Yang, Z. et al. Correlation of an epigenetic mitotic clock with cancer risk. Genome Biol. 17, 205 (2016).

20. Zhou, W. et al. DNA methylation loss in late-replicating domains is linked to mitotic cell division. Nat. Genet. 50, 591–602 (2018).

21. Teschendorff, A. E. A comparison of epigenetic mitotic-like clocks for cancer risk prediction. Genome Med. 12, 56 (2020).

22. Minteer, C. J. et al. Revisiting the bad luck hypothesis: Cancer risk and aging are linked to replication-driven changes to the epigenome. http://biorxiv.org/lookup/doi/10.1101/2022.09.14.507975 (2022) doi:10.1101/2022.09.14.507975.

23. Trapp, A., Kerepesi, C. & Gladyshev, V. N. Profiling epigenetic age in single cells. Nat. Aging 1, 1189–1201 (2021).

24. BIOS consortium et al. Age-related accrual of methylomic variability is linked to fundamental ageing mechanisms. Genome Biol. 17, 191 (2016).

25. Teschendorff, A. E. et al. Age-dependent DNA methylation of genes that are suppressed in stem cells is a hallmark of cancer. Genome Res. 20, 440–446 (2010).

26. Rakyan, V. K. et al. Human aging-associated DNA hypermethylation occurs preferentially at bivalent chromatin domains. Genome Res. 20, 434–439 (2010).

27. Feil, R. & Fraga, M. F. Epigenetics and the environment: emerging patterns and implications. Nat. Rev. Genet. 13, 97–109 (2012).

28. Nejman, D. et al. Molecular Rules Governing De Novo Methylation in Cancer. Cancer Res. 74, 1475–1483 (2014).

29. Levy, O. et al. Age-related loss of gene-to-gene transcriptional coordination among single cells. Nat. Metab. 2, 1305–1315 (2020).

30. Amit, G., Vaknin Ben Porath, D., Levy, O., Hamdi, O. & Bashan, A. Global coordination level in single-cell transcriptomic data. Sci. Rep. 12, 7547 (2022).

31. Horvath, S. & Raj, K. DNA methylation-based biomarkers and the epigenetic clock theory of ageing. Nat. Rev. Genet. 19, 371–384 (2018).

32. Hayflick, L. Entropy Explains Aging, Genetic Determinism Explains Longevity, and Undefined Terminology Explains Misunderstanding Both. PLoS Genet. 3, e220 (2007).

33. Lipsitz, L. A. Loss of ‘Complexity’ and Aging: Potential Applications of Fractals and Chaos Theory to Senescence. JAMA 267, 1806 (1992).

34. Gladyshev, V. N. et al. Molecular damage in aging. Nat. Aging 1, 1096–1106 (2021).

35. Yanai, S. & Endo, S. Functional Aging in Male C57BL/6J Mice Across the Life-Span: A Systematic Behavioral Analysis of Motor, Emotional, and Memory Function to Define an Aging Phenotype. Front. Aging Neurosci. 13, 697621 (2021).

36. Petr, M. A. et al. A cross-sectional study of functional and metabolic changes during aging through the lifespan in male mice. eLife 10, e62952 (2021).

37. Wang, S., Lai, X., Deng, Y. & Song, Y. Correlation between mouse age and human age in anti-tumor research: Significance and method establishment. Life Sci. 242, 117242 (2020).

38. Thomas, J. et al. Running the full human developmental clock in interspecies chimeras using alternative human stem cells with expanded embryonic potential. Npj Regen. Med. 6, 25 (2021).

39. Pyrkov, T. V. et al. Longitudinal analysis of blood markers reveals progressive loss of resilience and predicts human lifespan limit. Nat. Commun. 12, 2765 (2021).

40. Kerepesi, C., Zhang, B., Lee, S.-G., Trapp, A. & Gladyshev, V. N. Epigenetic clocks reveal a rejuvenation event during embryogenesis followed by aging. Sci. Adv. 7, eabg6082 (2021).

41. Sziráki, A., Tyshkovskiy, A. & Gladyshev, V. N. Global remodeling of the mouse DNA methylome during aging and in response to calorie restriction. Aging Cell 17, e12738 (2018).

42. Ying, K. et al. Causal Epigenetic Age Uncouples Damage and Adaptation. http://biorxiv.org/lookup/doi/10.1101/2022.10.07.511382 (2022) doi:10.1101/2022.10.07.511382.

43. Clemens, Z. et al. The biphasic and age-dependent impact of klotho on hallmarks of aging and skeletal muscle function. eLife 10, e61138 (2021).

44. Menichetti, G., Bianconi, G., Castellani, G., Giampieri, E. & Remondini, D. Multiscale characterization of ageing and cancer progression by a novel network entropy measure. Mol. Biosyst. 11, 1824–1831 (2015).

45. Sivakumar, S., LeFebre, R. W., Menichetti, G., Mugler, A. & Ambrosio, F. The fidelity of genetic information transfer with aging segregates according to biological processes. http://biorxiv.org/lookup/doi/10.1101/2022.07.18.500243 (2022) doi:10.1101/2022.07.18.500243.

46. Gladyshev, V. N. Aging: progressive decline in fitness due to the rising deleteriome adjusted by genetic, environmental, and stochastic processes. Aging Cell 15, 594–602 (2016).

47. Angermueller, C. et al. Parallel single-cell sequencing links transcriptional and epigenetic heterogeneity. Nat. Methods 13, 229–232 (2016).

48. Argelaguet, R. et al. Multi-omics profiling of mouse gastrulation at single-cell resolution. Nature 576, 487–491 (2019).

49. Hernando-Herraez, I. et al. Ageing affects DNA methylation drift and transcriptional cell-to-cell variability in mouse muscle stem cells. Nat. Commun. 10, 4361 (2019).

50. Cooper, G. M. et al. Distribution and intensity of constraint in mammalian genomic sequence. Genome Res. 15, 901–913 (2005).

51. Siepel, A., Pollard, K. S. & Haussler, D. New Methods for Detecting Lineage-Specific Selection. in Research in Computational Molecular Biology (eds. Apostolico, A., Guerra, C., Istrail, S., Pevzner, P. A. & Waterman, M.) vol. 3909 190–205 (Springer Berlin Heidelberg, 2006).

52. Yang, Z. Maximum likelihood phylogenetic estimation from DNA sequences with variable rates over sites: Approximate methods. J. Mol. Evol. 39, 306–314 (1994).

53. Thomas, J. W. et al. Comparative analyses of multi-species sequences from targeted genomic regions. Nature 424, 788–793 (2003).

54. Siepel, A. & Haussler, D. Computational identification of evolutionarily conserved exons. in Proceedings of the eighth annual international conference on Computational molecular biology - RECOMB ‘04 177–186 (ACM Press, 2004). doi:10.1145/974614.974638.

55. Siepel, A. et al. Evolutionarily conserved elements in vertebrate, insect, worm, and yeast genomes. Genome Res. 15, 1034–1050 (2005).

56. Felsenstein, J. & Churchill, G. A. A Hidden Markov Model approach to variation among sites in rate of evolution. Mol. Biol. Evol. 13, 93–104 (1996).

57. Lonsdale, J. et al. The Genotype-Tissue Expression (GTEx) project. Nat. Genet. 45, 580–585 (2013).

58. Battram, T. et al. The EWAS Catalog: a database of epigenome-wide association studies. Wellcome Open Res. 7, 41 (2022).

59. Liao, Y., Wang, J., Jaehnig, E. J., Shi, Z. & Zhang, B. WebGestalt 2019: gene set analysis toolkit with revamped UIs and APIs. Nucleic Acids Res. 47, W199–W205 (2019).

60. Cavalcante, R. G. & Sartor, M. A. annotatr: genomic regions in context. Bioinformatics 33, 2381–2383 (2017).

61. Fang, Y. et al. DNA methylation entropy is associated with DNA sequence features and developmental epigenetic divergence. Nucleic Acids Res. 51, 2046–2065 (2023).

62. Luo, C., Hajkova, P. & Ecker, J. R. Dynamic DNA methylation: In the right place at the right time. Science 361, 1336–1340 (2018).

63. Zhang, B. et al. Multi-omic rejuvenation and lifespan extension upon exposure to youthful circulation. http://biorxiv.org/lookup/doi/10.1101/2021.11.11.468258 (2021) doi:10.1101/2021.11.11.468258.

64. Olova, N., Simpson, D. J., Marioni, R. E. & Chandra, T. Partial reprogramming induces a steady decline in epigenetic age before loss of somatic identity. Aging Cell 18, e12877 (2019).

65. Gill, D. et al. Multi-omic rejuvenation of human cells by maturation phase transient reprogramming. eLife 11, e71624 (2022).

66. MAMMALIAN METHYLATION CONSORTIUM et al. Universal DNA methylation age across mammalian tissues. http://biorxiv.org/lookup/doi/10.1101/2021.01.18.426733 (2021) doi:10.1101/2021.01.18.426733.

67. Zhang, B. et al. Epigenetic profiling and incidence of disrupted development point to gastrulation as aging ground zero in Xenopus laevis. http://biorxiv.org/lookup/doi/10.1101/2022.08.02.502559 (2022) doi:10.1101/2022.08.02.502559.

68. Minteer, C. et al. Tick tock, tick tock: Mouse culture and tissue aging captured by an epigenetic clock. Aging Cell 21, (2022).

69. Kabacik, S. et al. The relationship between epigenetic age and the hallmarks of ageing in human cells. Nat. Aging (2022) doi:10.1038/s43587-022-00220-0.

70. Kerepesi, C. et al. Epigenetic aging of the demographically non-aging naked mole-rat. Nat. Commun. 13, 355 (2022).

71. Horvath, S. et al. DNA methylation clocks tick in naked mole rats but queens age more slowly than nonbreeders. Nat. Aging 2, 46–59 (2022).

72. Battich, N., Stoeger, T. & Pelkmans, L. Control of Transcript Variability in Single Mammalian Cells. Cell 163, 1596–1610 (2015).

73. Raj, A. & Van Oudenaarden, A. Nature, Nurture, or Chance: Stochastic Gene Expression and Its Consequences. Cell 135, 216–226 (2008).

74. Virtanen, P. et al. SciPy 1.0: fundamental algorithms for scientific computing in Python. Nat. Methods 17, 261–272 (2020).

75. Teschendorff, A. E., Breeze, C. E., Zheng, S. C. & Beck, S. A comparison of reference-based algorithms for correcting cell-type heterogeneity in Epigenome-Wide Association Studies. BMC Bioinformatics 18, 105 (2017).

76. Zheng, S. C. et al. A novel cell-type deconvolution algorithm reveals substantial contamination by immune cells in saliva, buccal and cervix. Epigenomics 10, 925–940 (2018).

77. Zheng, S. C. et al. EpiDISH web server: Epigenetic Dissection of Intra-Sample-Heterogeneity with online GUI. Bioinformatics 36, 1950–1951 (2020).

78. Kent, W. J. et al. The Human Genome Browser at UCSC. Genome Res. 12, 996–1006 (2002).

79. Menees, K. B. et al. Sex- and age-dependent alterations of splenic immune cell profile and NK cell phenotypes and function in C57BL/6J mice. Immun. Ageing 18, 3 (2021).

80. Sloan, C. A. et al. ENCODE data at the ENCODE portal. Nucleic Acids Res. 44, D726–D732 (2016).

